# Microchip based microrheology via Acoustic Force Spectroscopy shows that endothelial cell mechanics follows a fractional viscoelastic model

**DOI:** 10.1101/2020.07.02.185330

**Authors:** Alfred Nguyen, Matthias Brandt, Timo Betz

**Affiliations:** Institute for Cell Biology, University of Münster, Münster, Germany

## Abstract

Active microrheology is one of the main methods to determine the mechanical properties of cells and tissue, and the modelling of the viscoelastic properties of cells and tissue is under heavy debate with many competing approaches. Most experimental methods of active microrheology such as optical tweezers or atomic force microscopy based approaches rely on single cell measurements, and thus suffer from a low throughput. Here, we present a novel method for cell based microrheology using acoustic forces which allows multiplexed measurements of several cells in parallel. Acoustic Force Spectroscopy (AFS) is used to generate multi-oscillatory forces in the range of pN-nN on particles attached to primary human umbilical vein endothelial cells (HUVEC) cultivated inside a microfluidic chip. While the AFS was introduced as a single-molecule technique to measure mechanochemical properties of biomolecules, we exploit the AFS to measure the dynamic viscoelastic properties of cells exposed to different conditions, such as flow shear stresses or drug injections. By controlling the force and measuring the position of the particle, the complex shear modulus *G*\*(*ω*) can be measured continuously over several hours. The resulting power-law shear moduli are consistent with fractional viscoelastic models. In our experiments we confirm a decrease in shear modulus after perturbing the actin cytoskeleton via cytochalasin B. This effect was reversible after washing out the drug. Although these measurements are possible, we provide critical information regarding the AFS as a measurement tool showing its capabilities and limitations. A key result is that for performing viscoelastic measurements with the AFS, a thorough calibration and careful data analysis is crucial, for which we provide protocols and guidelines.

## Introduction

The mechanical properties of cells and tissues are closely related to their biological function, and defects in stiffness and viscosity have been related to several malfunctions and diseases. ^1–4^ An example of cells highly exposed to variable mechanical forces are endothelial cells (ECs) which make up the inner wall of blood vessels. They are constantly exposed to variable shear stresses originating from the blood flow. These forces are known to be important regulators for proper EC function. ^5,6^ A highly relevant example are EC stiffness dependent biochemical processes which can promote vascular diseases, such as atherosclerosis. ^7,8^ Changes in stiffness between healthy and diseased cells are also often found in other cell types, such as cancer. ^3,9^ The cell mechanical response to different exposures, such as biochemical stimuli, is an important characterization to better understand the processes in healthy and malfunctional cells. Therefore, the study of cell mechanical properties becomes of great importance, and has triggered many studies in the past decades. Despite the large interest in quantitative descriptions of cellular viscoelastic properties, a clear model description is not yet available. ^10^ While in the early days of cell mechanics, classical analogy models for viscoelasticity using springs and dash-pots, like the Maxwell, the Kelvin-Voigt or the standard solid viscoelastic model were used ^11,12^, it became clear that cells can be best described by power-law rheology models. ^13^ Recently, a fractional viscoelastic element has turned out to be a highly flexible approach to describe the different experimental data measured in microrheology. However, a single fractional element is often insufficient to describe the experimentally found differences in the power-law exponents of the storage and the loss modulus. ^14–17^ One key finding of the work presented here is that microrheology of ECs can be well modelled by the generalized Kelvin-Voigt model comprised of two fractional elements in parallel.

Microrheology is a major method to determine cell mechanical properties. Currently, there are different microrheology techniques of adherent cells available, such as optical tweezers ^18–21^, atomic force microscopy ^22–25^ and magnetic twisting cytometry. ^12,26–28^ Each of them have their own advantages and drawbacks. Optical tweezers (OTs) use a highly focused laser to exert forces on particles. While the force range is typically limited to a few hundred pN, OTs have the advantage of easy access to sealed samples, an accurate control of the trap location in 3D, and a high temporal and spatial resolution. The disadvantages are that only single measurements are possible, and only small forces can be applied to avoid local heating at the focus of the laser trap which may affect the measured properties of cells. ^29^ The throughput can be increased by using holographic optical tweezers (HOTs) at the cost of the absolute position detection ^30,31^ or time-shared optical tweezers (TS-OTs) at the cost of time resolution and possible oscillation artifacts. ^32,33^ Another technique for microrheology is the atomic force microscopy (AFM). ^22,34,35^ AFMs use a cantilever with a tip with different shape designs depending on the experiment to exert forces on a substrate that it is approaching. While a large force range of pN to a few nN can be generated and AFMs are also capable of imaging the substrate, they need open chambers and only one measurement can be performed at a time, again limiting throughput. Besides these main methods, further techniques have been developed, such as magnetic tweezers ^36^, stretching rheometer ^37^, optical stretcher ^38^, microfluidic approaches ^39^ and micropipette aspiration. ^40^

With the exception of the magnetic twisting cytometry, these techniques typically only measure one particle or location at a time. More importantly, they usually do not easily allow a measurement during a fluid flow as open chambers are required or the shear forces of the fluid flow would be too high. In this work, we present a novel method for microrheology based on acoustic forces using the recently introduced technique of Acoustic Force Spectroscopy (AFS) which can overcome these drawbacks. The AFS is initially designed to measure mechanochemical properties of biomolecules in a high-throughput manner. ^41,42^ Recently, the AFS has been used to stretch red blood cells under a constant force on attached particles to measure their static spring constant. ^43^ In our work, we apply a custom, time-varying, oscillatory force on particles that are attached on a monolayer of primary human umbilical vein endothelial cells (HUVEC). With this new method the results using the AFS can be compared with other techniques, such as AFMs. The aim of this work is to implement the AFS for microrheology and to determine the viscoelastic properties of HUVEC monolayers under variable conditions, while testing recent viscoelastic material models like different versions of fractional elements.

## Materials and Methods

### Acoustic Force Spectroscopy

The Acoustic Force Spectroscopy (AFS; LUMICKS B.V, Netherlands) equipment consists of a Generation 2 AFS module containing a function generator and a temperature controller. Visualization is achieved by an inverted microscope setup with a motorized 10 × objective (Nikon, Plan Fluor) for a nanometer-precise *z*-translation, an LED and a uEye camera (UI324xML-M) capable of imaging 1280 × 1024 pixels corresponding to a 678.40 *μ*m×542.72 *μ*m field of view with a sampling frequency of 59 Hz. The Generation 2 chips consist of a flow chamber with a fill volume of about 6 *μ*L and a piezoelectric element (piezo) on the top of the glass chip to transduce the acoustic waves. ^41^ Acoustic forces are applied on 10 *μ*m diameter polystyrene beads (micromer, micromod Partikeltechnologie GmbH). During the experiment the beads are tracked by image recording using the LabVIEW software provided by the manufacturer (LUMICKS B.V) which we have modified for our needs. The *z*-position of each bead is determined using a predefined lookup-table (LUT) ranging from 0 nm to 20000 nm in 100 nm steps according to van Loenhout *et al*. ^44^ Real time image processing allows a live-view of the bead position. The software also generates the voltage signals for the piezo to apply the acoustic pressure which results in a force on the beads. The modified software can generate signals to exert custom, time-varying force profiles, such as force oscillations, and is compatible with the self-written MATLAB (The MathWorks, Inc.) analysis program *Kitsune* (available on github https://github.com/A141-User/Acoustic-Kitsune) enabling a live-view of the recorded data which can be analyzed afterwards.

### Stokes Force Calibration of AFS chips

As the software generates voltage signals, a force calibra-tion is required. Stokes Force Calibration (SFC) yields the conversion factor from voltage to force for the 10 *μ*m-diameter polystyrene beads. Fluorescent (FluoSpheres™, F8834, Thermo Fisher Scientific) and carboxylated beads (micromer, micromod Partikeltechnologie GmbH) are used in culture medium (ECGM/M199 mixture) and water, respectively. Beads are suspended to a concentration of approximately 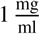 and injected in the chip. This concentration avoids bead clustering and high bead density which would prevent tracking. Temperature is set using the LabVIEW software and the feedback-controlled temperature sensor mounted on the chip. For each bead inside the field of view (FoV) a look-up-table (LUT) between each bead’s radial intensity profile and its *z*-position is generated. To displace the beads from the chip surface at least three different acoustic amplitudes (in %) are applied for 1 s. The time between applying these amplitudes is 15 s and 11 s for the SFC in ECGM/M199 and in water, respectively. Typically, about 25 beads at random positions are measured simultaneously inside a field of view (FoV). This procedure is repeated after flushing new beads in the FoV. As the conversion factor depends on the position in the FoV, a spatial calibration map is generated by measuring at least more than 1000 beads with well-distributed data points. In case of problematic calibration, such as beads binding unspecifically to the glass surface or tracking errors due to bead clustering, these beads are filtered out before generating the spatial map. A protocol of the SFC and more details on data analysis of the SFC are given in the **SI**.

### Determination of the bead immersion half-angle

Human umbilical vein endothelial cells (HUVECs) are seeded overnight on a 35 mm diameter glass bottom dish (Greiner Bio CELLVIEW Cell Culture Dish) until they form a confluent monolayer. The glass surface has been coated with 68 *μ*g/ml collagen type I from rat tail (Corning) for 1 h. 10 *μ*m diameter collagen coated polystyrene beads (micromer, micromod Partikeltechnologie GmbH) are added on top of the HUVEC monolayer and are allowed to sink for 30 min. During the same time period the cell membrane is stained using a plasma membrane dye (CellMask™ Deep Red, ThermoFisher Scientific) at 0.5 *μ*M concentration. 3D-stacks of 90 Z-planes à 0.2 *μ*m distance for five different fields of view are acquired with a scientific CMOS camera (Prime BSI Photometrics) using spinning disk confocal microscopy (Nikon Eclipse Ti-E, equipped with CSUW1 Yokogawa SD head) using a 60X Plan Apo water-immersion objective with a numerical aperture of 1.27 and an excitation laser of 640 nm wavelength. Using a custom MATLAB code the cell surface was determined by locally fitting polynomials to the fluorescence intensity data. For a total of 43 beads the diameter *d* (see Fig. 1B) of the bead-cell contact circle was determined averaging over 4 different contact points chosen by eye. Knowing the bead radius *R* the immersion half-angle *0* was calculated as

**Figure 1.**
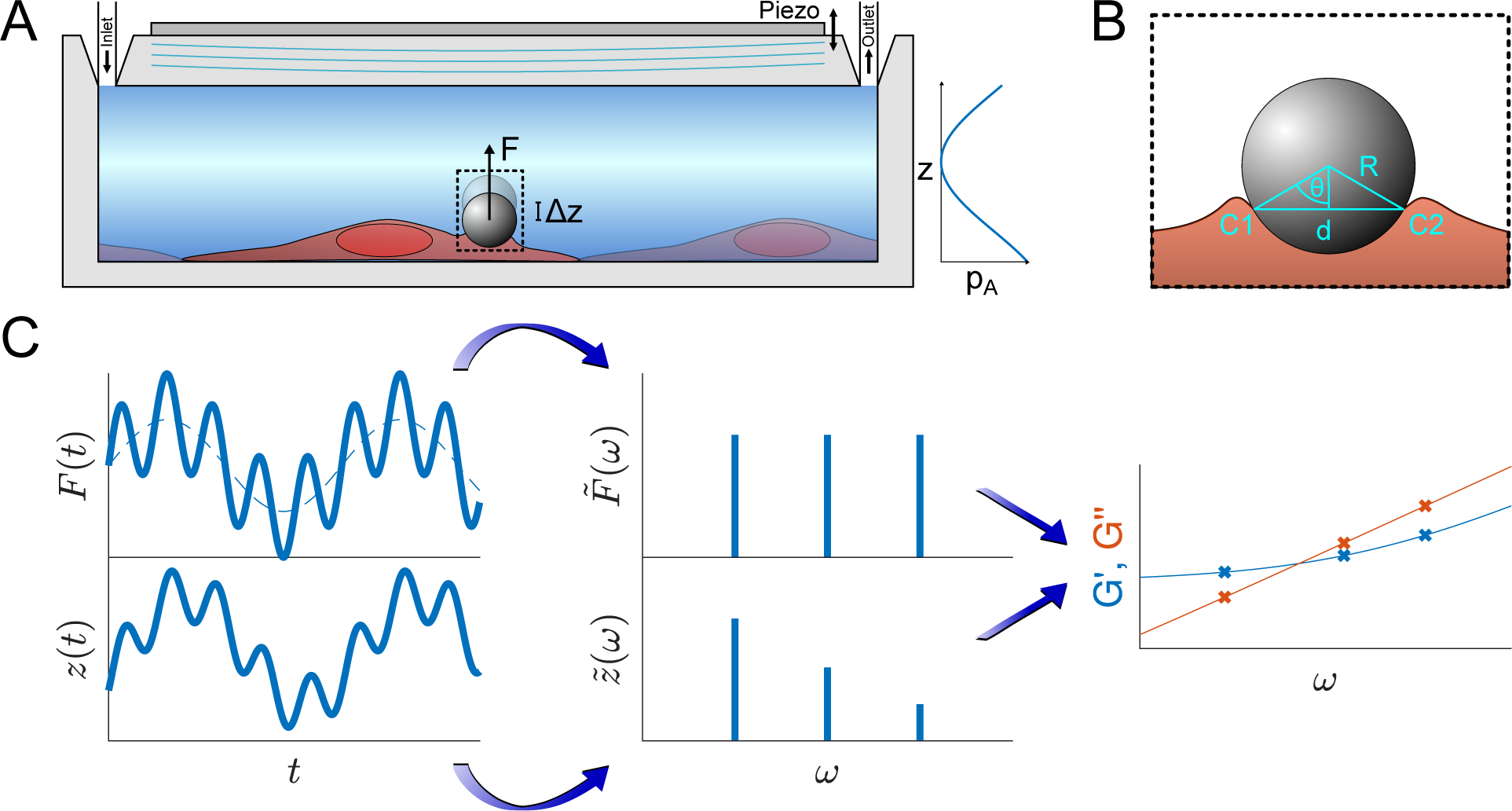
Principle of the microrheological experiment using the AFS. (A) Schematic cross-section of the AFS showing the fluid chamber with a cell monolayer. The oscillating piezoelectric element generates a standing pressure wave *p*_*α*_ inside the fluid chamber resulting in a force in the *z*-direction on a small bead attached on the cell. (B) Illustration of a partially immersed bead on a cell surface. Bead and cell surface come in contact at points C1 and C2. The distance between C1 and C2 represents the diameter *d* of the bead-cell contact circle, *R* denotes the bead radius and *θ* the immersion half-angle. (C) Analysis of the multi-oscillation microrheology. The force is modulated with multiple superimposed frequencies and the measured *z*-position follows with the same frequencies. The Fourier transform 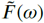 and 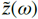 of the force and the *z*-position is calculated. The resulting complex shear modulus is then obtained with Eq. 7.

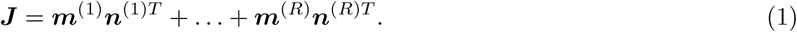

### Microrheological experiment

The principles of the microrheological experiment are shown in Fig. 1. The chip with the HUVEC monolayer is placed on the AFS microscope setup and temperature controlled to 36°C ± 1°C to avoid temperatures above 37°C given the uncertainty of the heating system. The connected syringe pump is kept at a flow rate of 1.66 *μ*l/min. Through an injection site 10 *μ*m collagen coated polystyrene beads (micromer, micromod Partikeltechnologie GmbH) suspended in CO_2_-*charged medium* are inserted using a syringe. After closing the outlet valve to prevent flow during the bead incubation time (about 5 min) the valve is reopened and the resulting flow removes unattached beads. This process is repeated until a desired amount of beads has attached to the monolayer inside the calibrated field of view. Then, the flow is stopped for 5 min and the temperature is de-creased to approximately 33°C as the acoustic force application increases the temperature inside the chip. After generating the look-up-table (LUT) for each bead in the FoV, the acoustic forces inside the AFS chip are applied, pulling the beads upwards, away from the surface. For the multi-frequency oscillatory (*mOsc*) force the frequencies 0.1 Hz, 0.5 Hz and 1.5 Hz are applied simultaneously. Continuous mOsc forces are exerted on the beads attached on the cells of the monolayer for about 3700 s using the following protocol for drug treatment and control conditions:

a. for 1000 s a flow rate 1.66 *μ*l/min of CO_2_-*charged medium* is applied to mimic the culture conditions;
b. change of medium containing syringe to apply control (CO_2_-*charged medium*) or treatment (1 *μ*g/mL cytochalasin B, Sigma-Aldrich) medium for 1000s at a flow rate 30 *μ*l/min. As the volume to be filled is about 300 *μ*l the new conditioned medium arrives at the chamber after 600 s;
c. finally, the flow rate is decreased to the initial 1.66 *μ*l/min for 1700 s. The syringe exchange takes less than 100 s, and during this change the flow rate is initially lower and then higher when adjusting the pusher block of the syringe pump to the plunger of the syringe.

After this protocol the mOsc force is stopped and the flow rate is shortly increased to 30 *μ*l/min to remove detached beads and then the temperature is set back to 36°C and the flow rate to 1.66 *μ*l/min.

To confirm cell recovery after cytochalasin B (cyto B) treatment, the mOsc force measurement is not stopped after c), but the syringes are exchanged back to the CO_2_-*charged medium* at a flow rate of 30 *μ*l/min for 1100 s. Then, the flow is decreased again to 1.66 *μ*l/min until the end of the measurement.

### Cell culture

Human umbilical vein endothelial cells (HUVECs) are kindly provided by Prof. Dr. V. Gerke (Institute of Medical Biochemistry, University of Münster, Münster, Germany) and are obtained according to the protocol described in Jaffe *et al*. ^45,46^ HUVECs are cultivated in a 60 mm × 15 mm culture dish (Corning Cell Culture Dish, Corning CellBIND Surface, non-pyrogenic, polystyrene) at 37°C and 5 % CO_2_ in a humidified incubator. Culture medium consists of a 50/50 mixture of two medium solutions; namely, Endothelial Cell Growth Medium 2 (ECGM; PromoCell, Germany) supplemented with the SupplementMix for ECGM 2 (PromoCell), and Medium 199 Earle’s (F0615, Biochrom GmbH, Germany or M2154, Sigma-Aldrich) with *2*.2 g/l NaHCO_3_, without L-glutamine, supplemented with 10 % fetal bovine serum (F7524, SigmaAldrich), 30 *μ*g/ml gentamicin (gibco), 0.015 *μ*g/ml amphotericin B (gibco) and 0.2 units/ml heparin (H3149, Sigma For flow experiments, CO_2_-*charged medium* was used, which refers to the described cell culture medium, which is pH-stabilized by equilibration in a 5 % CO_2_ atmosphere. HUVECs of passages 3-6 are split at 70 − 90 % confluence, and are discarded after passage 6. To allow cell spreading, the flow chamber of the AFS chip is coated with 68 *μ*g/ml collagen type I from rat tail (Corning) for at least 4 h at room temperature. HUVECs are seeded onto the flow chamber of the AFS chip at a density of about 2 × 10^6^ cells/ml before placed into a dry incubator at 37°C and 5 % CO_2_. For long term visualization of cells, the chip is mounted on the AFS equipment and heating is ensured by the temperature controller of the instrument. After > 2 h most of the cells have attached onto the surface. Using a syringe pump (AL-2000, World Precision Instruments) CO_2_-*charged medium* is pumped through the flow chamber at a flow rate of 500 *μ*l/min for *<* 1 min to flush out non-attached and possible dead cells. Then, the flow rate is set to 1.66 *μ*l/min to renew the medium during cultivation. For the monolayer experiments, HUVECs are grown inside the incubator for 70 − 100 h.

### Data analysis

#### Power-law rheology models

As we observe power-law behavior for the viscoelastic shear modulus, recent fractional models are considered to explain the data. A detailed explanation of the models is found in the SI. Briefly, we apply either a single fractional element model (SFE) or a Generalized Kelvin-Voigt model (GKV) that is based on a combination of fractional elements. ^47^ In the SFE model, the complex shear modulus is:

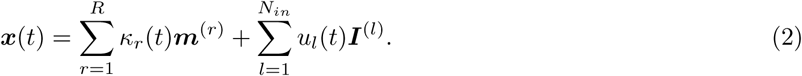

yielding a power-law behavior as often ^37,48^ used to describe linear viscoelastic materials. Important is that in the SFE, the power-law exponent for the real and the imaginary part are the same, which is not found in our results.

The second model is a combination of two SFEs resulting in the GKV model:

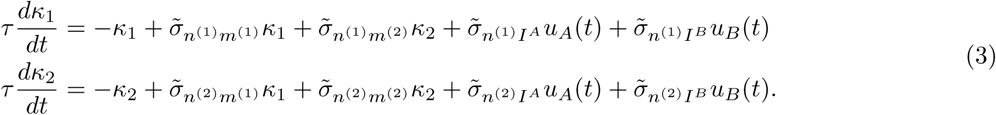

This is further simplified by setting *t*_0_ = 1 s that leads to 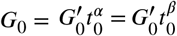 thus yielding the expression of the real and imaginary part as:

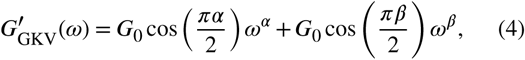

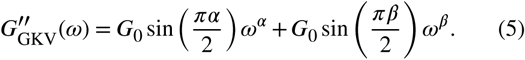

The simplification yields expressions (Eqs. 4, 5) with three independent parameters (*G*_0_, *α, β*).

This model gives a crossing of the real and imaginary part at the crossover frequency

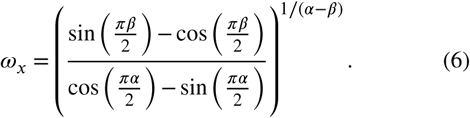

In the Supplementary Information (Fig. S.7) certain selected cases for different *α, β* are shown for the complex shear modulus obtained with the SFE, GKV and the Generalized Maxwell (GM) model.

To obtain the stress *σ* and the strain *ϵ* from the measured force and bead displacement *z* we follow the approach previously developed for an optical tweezers experiment. ^37,49^ The resulting complex modulus is calculated depending on the immersion of a bead inside the material characterized by an immersion half-angle *θ*, see Fig. 1B, i.e.

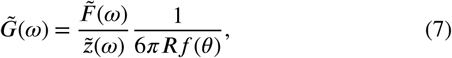

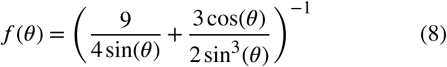

with [*f* (*θ*) *<* 1, ∀ *θ* ∈ (10°, 90°)], see Supp. Fig. S.9, which will be used for the analysis. For a bead immersed in an infinite medium ^37,50^, the complex modulus would be 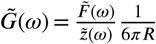 which is identical to the generalized Stokes-Einstein relation.

#### Data post-processing

The obtained data has been post-processed using our software *Kitsune* to obtain correct values for the viscoelasticity. The software filters for inter-particle errors in the tracking, corrects for drift in the long time measurements and includes the local distribution of the conversion factor. Details about the analysis are found in the SI. Briefly, the software corrects for the following three main problems:

a. During the constant medium flow, other particles can interfere with the tracking of the beads that will cause erroneous values in the (*x, y, z*)-position data. Typically, these erroneous values only appear for a short time, i.e. one or two data points, due to the relatively high flow rate compared to the sampling frequency. This problem is addressed by replacing the erroneous value with the median value of data points temporally close to it.
b. The long measurement time may introduce mechanical drifts or allows drifts caused by active, but slow, cell movements over time. The drift is corrected by obtaining the drift profile with segmented, continuous fits of polynomial functions of second order and subtracting them from the data. This correction can be performed because the cell movement and activity themselves are not of interest in this case.
c. The lateral inhomogeneity of the force distribution inside the field of view will cause a wrong assignment of the actual force acting on the beads. This can be corrected by assigning a conversion factor of the amplitude to force for each bead according to their current lateral position with the obtained spatial map from the SFC, as described in the next section.

## Results

A main result of this work is to provide a clear description for the calibration of the AFS, and to obtain relevant frequency dependent microrheology data for the ECs monolayer. For successful microrheology experiments we found that special care needs to be taken for the bead location, the resonance frequency at the precise temperature and the details of the medium used. To get from the forces to the shear modulus we also determine the bead indentation. Only when these points are well controlled, the microrheology experiments using the AFS yield reliable results.

### Stokes Force Calibration of AFS chips

Knowledge of the exerted acoustic force inside the AFS chip is essential to use the AFS as a measurement tool. In this section we provide a new calibration model with respect to previous approaches ^42^, and compare the different model with an independent measurement.

Literature reveals the expression of the acoustic radiation force in *z* acting on a spherical particle with radius *R*, compressibility *κ*_*p*_ and density *ρ*_*p*_ in a medium with compressibility *κ*_*m*_ and density *ρ*_*m*_ as ^42,51,52^

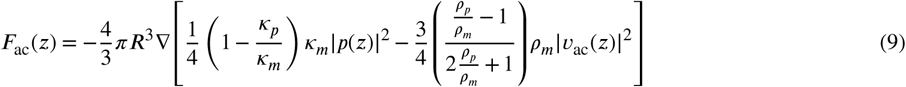

with the acoustic pressure *p*(*z*) and acoustic velocity *v*_ac_(*z*). Further details are given in the Supplementary Information (Eq. S.1). Sinusoidal voltage signals from the function generator are converted to acoustic pressure waves by the piezoelement. The acoustic intensity is controlled by an amplitude value in percentage [%] units. For the conversion from [%] units to force units, e.g. pN, a conversion factor *c* is determined by calibration. The calibration method provided by the manufacturer for *in situ* relies on beads tethered to small linear molecules, such as DNA, and exploits the Brownian motion of the particles. ^41^ However, this method is unsuitable for beads attached to cells, as the bead motion is dominated by the properties of the cell, and not of the applied force. Hence, for cell microrheological experiments an *in situ* calibration is currently not available. To overcome this, we use the *Stokes Force Calibration* (*SFC*) that is based on the drag force. Previous applications ^42^ of the Stokes forces applied a viscosity correction to the wall effects that is valid for motion close and parallel to a surface (Faxén’s law). As in AFS experiments the bead moves normal to the surface Faxén’s law is not suitable. Here, we present another viscosity correction factor derived for motion normal to the surface that leads to excellent agreement with the measurements.

When the acoustic force *F*_ac_ is applied on a bead suspended in medium, it is in a force equilibrium

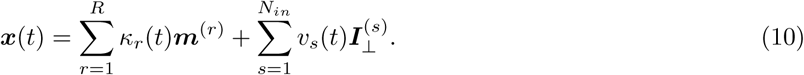

with the effective gravitational force *F*_eff. grav._ and the Stokes drag force *F*_Stokes_. Due to the low Reynolds number of the system inertia can be neglected. For a spherical bead the effective gravitational force is given by

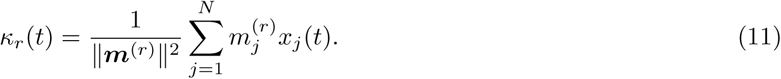

with the gravitational constant 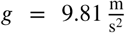 and the compressibility-corrected temperature-dependent density of water, see Appendix (Eq. A.2).

The Stokes drag force is given by

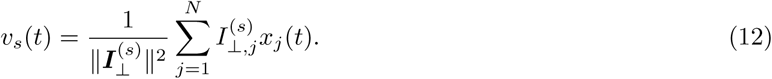

with the bead velocity *v*, viscosity *η* and the viscosity correction factor *λ*. As the viscosity is temperature-dependent it can be approximated for water with a modified Andrade equation^53^, see Appendix (Eq. A.3). The viscosity of cell culture medium containing serum is approximated to be the same as water. This is justified by measurements showing that the presence of serum only increases the viscosity by about 5 %. ^54^ Since the bead is very close to the surface, the *z*-position-dependent viscosity correction *λ* becomes of great importance. In the case of measurements with the AFS where the bead translation is perpendicular to the surface, Faxén’s law for *λ* (see Appendix, Eq. A.5) should not be used because it is a correction for a translation parallel to the surface and would yield wrong calibration results as shown later (Fig. 2). Instead, the viscosity correction for perpendicular translation as given by Brenner is used ^55^:

**Figure 2.**
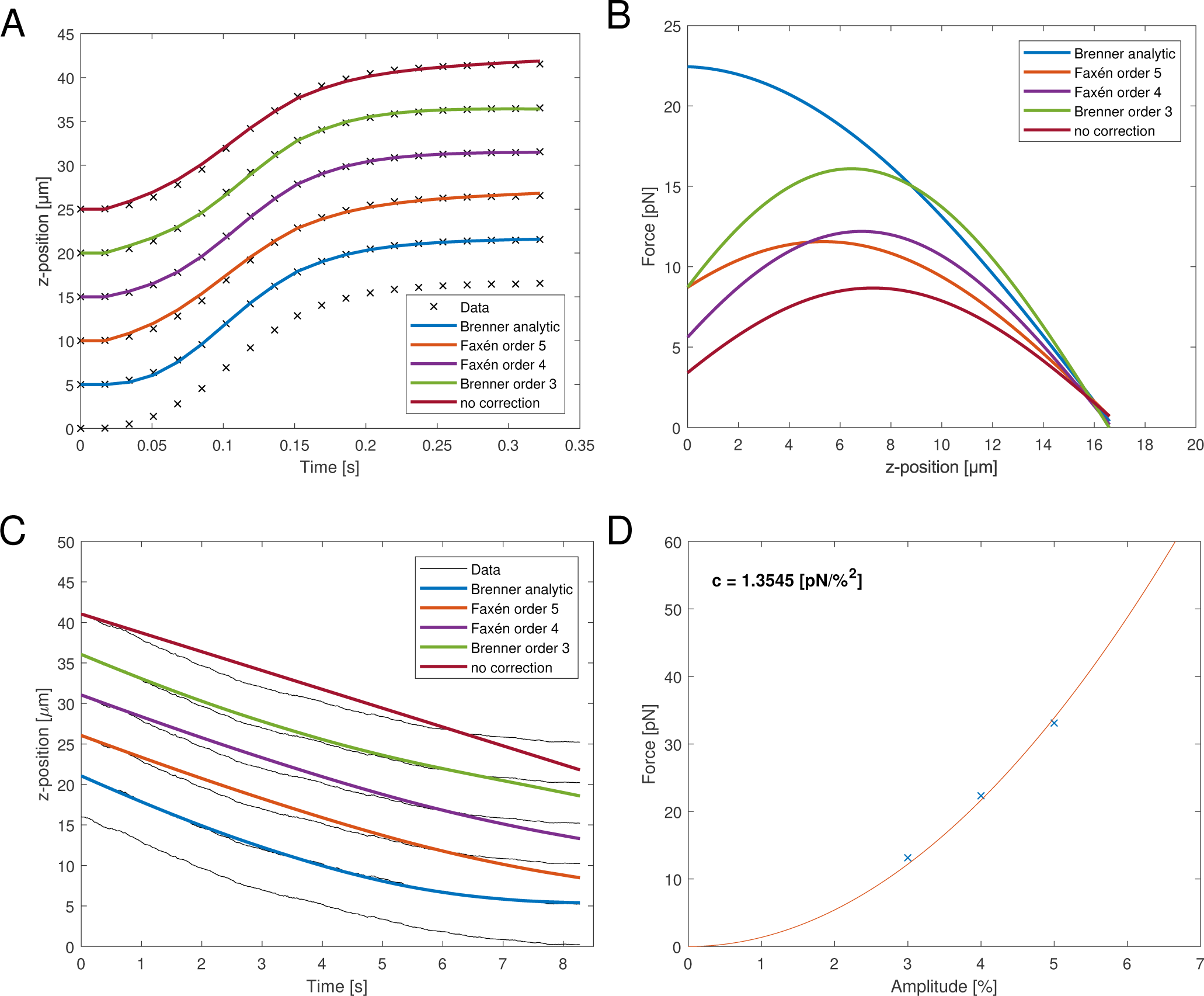
(A) Representative, measured *z*-position data (black cross) obtained after SFC measurement and the fits by integrating Eq. 16 with the different viscosity corrections. The goodness of fit is *R*^2^ > 0.9971 for all shown fits. For clarity, the different fits and the data were shifted by 5 *μ*m from each other. (B) Resulting force profile obtained from the fit parameters with Eq. 14 of the positional data from (A) with the different viscosity corrections. The fit parameter values are shown in Table 1. The applied amplitude was 4 %. (C) Representative, measured *z*-position data (black line) of a falling bead after the SFC and the fits with different viscosity corrections. The fits and data were shifted by 5 *μ*m from each other for clarity. The density of the bead was used as the fit parameter and the values are shown in Table 1. (D) Parabolic relationship between the acoustic force and the applied amplitude. The force was calculated with the viscosity correction with Eq. 13 and the force at *z* = 1 *μ*m was used to obtain the conversion factor *c*.

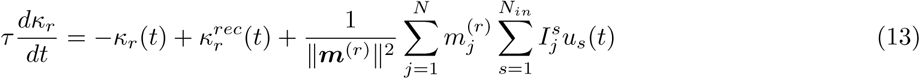

**Table 1.**
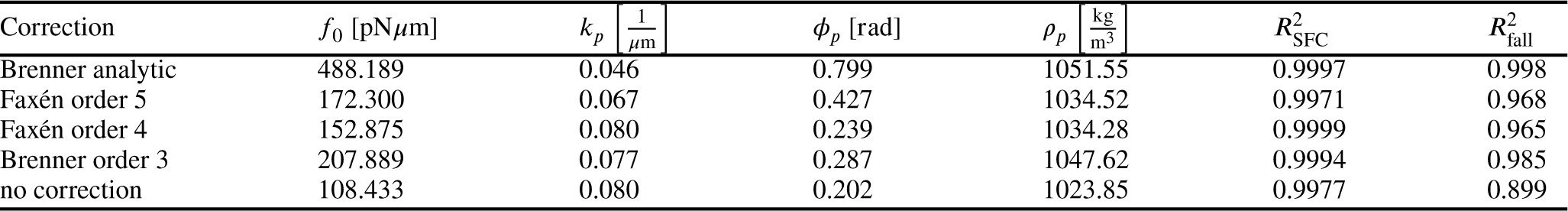
Values of the fit parameters to calculate the force profile and the goodness of fit obtained by the SFC and the bead fall shown in Fig. 2A, 2C.

| Correction | $f_0$ [pNμm] | $k_p$ [$\frac{1}{\mu$m}] | $\phi_p$ [rad] | $\rho_p$ [$\frac{\text{kg}}{\text{m}^3}$] | $R_{\text{SFC}}^2$ | $R_{\text{fall}}^2$ |
| --- | --- | --- | --- | --- | --- | --- |
| Brenner analytic | 488.189 | 0.046 | 0.799 | 1051.55 | 0.9997 | 0.998 |
| Faxén order 5 | 172.300 | 0.067 | 0.427 | 1034.52 | 0.9971 | 0.968 |
| Faxén order 4 | 152.875 | 0.080 | 0.239 | 1034.28 | 0.9999 | 0.965 |
| Brenner order 3 | 207.889 | 0.077 | 0.287 | 1047.62 | 0.9994 | 0.985 |
| no correction | 108.433 | 0.080 | 0.202 | 1023.85 | 0.9977 | 0.899 |

with 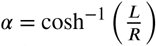 and *L* the distance of the center of the bead to the surface, i. e. *L* = *z* + *R*. For the numerical computation the sum goes from 1 to 100.

The acoustic force in Eq. 9 can be further simplified by recalling that polystyrene in water or culture medium is near-neutral buoyant, so the second term containing the acoustic velocity is small compared to the first term, and is hence neglected. However, the gravitational force is still considered in the analysis. The applied pressure has a sinusoidal form, e.g. *p*(*z*) = *p*_0_ cos(*k*_*p*_*z* + *ϕ*_*p*_) with the pressure amplitude *p*_0_, wave number K_*p*_ and phase *ϕ*_*p*_. Thus, the acoustic force can be approximated by

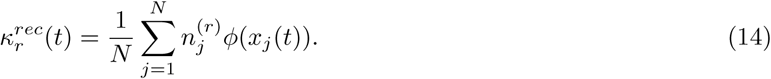

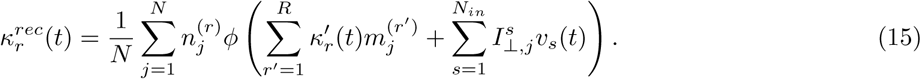

With Eq. 10 and Eq. 14 the *z*-position can be obtained by numerically integrating the velocity

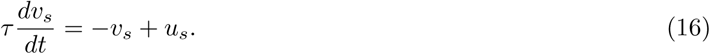

For the calibration, the fit parameters *f*_0_, *k*_*p*_ and *ϕ*_*p*_ are optimized until the predicted *z*_num_ and measured *z*_meas_ positions during the force application match best. Thus, neither the knowledge of the compressibility of the particle nor of the medium is required for the SFC.

To demonstrate the effect of the viscosity corrections of Faxén’s law and Brenner’s solution (see Appendix: Eqs. A.5, A.6) are compared for different orders in Figure 2. We compare the correction for parallel translation Faxén’s law at different orders (Faxén: order 4, Faxén: order 5), and a third order approximation (Brenner: order 3) as well as the analytic Brenner correction (Brenner: analytic) for perpendicular translation. Figure 2A shows the data and the different fits of the *z*-position during the acoustic force application. At first sight, the different viscosity corrections all yield a *z*-position that fits the measured data with a high fit quality, suggesting that all approximations are equally good (see Table 1). However, when using the obtained fit parameters to calculate the acoustic force profile in the *z*-dimension with Eq. 14 it becomes evident that the predicted forces are drastically different (Fig. 2B), especially in the relevant region close to the surface (small *z* values). The main difference are the wave number and phase. These fix the *z*-position of the maximum force. For the analytic Brenner correction Eq. 13 the maximum is at the surface of the chip, as expected, which means that it can be approximated as constant close to the chip surface. In contrast, for the other correction factors the maximum is at a larger distance of *z* > 5 *μ*m from the surface. Furthermore, the absolute value of the prefactor *f*_0_ which yields information of the force varies by almost a factor of five depending on the analysis (see Table 1).

This result shows that a good fit for the displacement curve upon force application is not sufficient to get a reliable calibration. To check the different approximations independently we validated the SFC calibration by observing the bead falling from a height *z* > 0 *μ*m after turning off the acoustic force. The only relevant forces are now the gravitational force and the Stokes drag force. Here, only the density of the bead *ρ*_*p*_ is used as a fit parameter. Figure 2C shows that the analytic viscosity correction (Eq. 13) is indeed the suitable correction compared to the other corrections because the obtained *z*-position fits the measured data the best 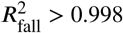 as well as the density of the bead is closest to the expected value of 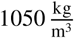, see Table 1. Now that the acoustic force at different amplitudes has been measured, a conversion factor can be obtained using the analytic viscosity correction. The relation between the force *F*_ac_ and the amplitude *V*_%_ is quadratic

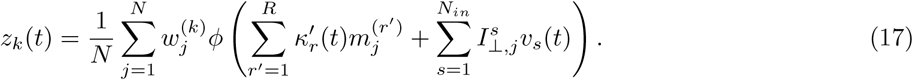

with the conversion factor *c* in 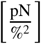 because the amplitude is directly proportional to the pressure. The conversion factor is obtained by fitting a polynomial function of second order (Eq.17) to at least two different, measured force values at a non-zero amplitude (*V*_%_ ≠ 0 %), see Fig. 2D. Indeed, the quadratic relation can be seen in the measured data. The force values are obtained from the force profiles at *z* = 1 *μ*m.

### Lateral force heterogeneity in the AFS chip

For an ideal system, i.e. an infinite surface without boundary and a perfectly perpendicularly aligned pressure wave, the forces on small particles are independent on the lateral position of the particles and the forces are only exerted in the *z*-direction. However, we observe a lateral translation of free beads when applying an acoustic force (see Supp. Fig. S.1). This indicates relevant lateral forces that cannot be neglected, and hints for a lateral position dependence of the force conversion factor. The spatial dimension of the flow chamber underneath the transparent piezo is shown in Fig. 3A, 3B. In order to create a spatial map representing the conversion factor *c* at different lateral positions, the SFC is performed on over 1000 randomly distributed beads inside the same field of view (FoV) shown in Fig. 3C. As shown in Fig. 3D the resulting force distribution in the chip is highly inhomogeneous, and varies up to a factor of six depending on the position of the bead. The most efficient force generation is not directly at the center of the chip in the short axis (*x*-position), but slightly shifted outwards. In fact, typically the change of the force is even very high close to the center; in the shown FoV (Fig. 3C) the center of the flow chamber is at around 100 *μ*m. Only about a quarter of the short axis of the flow chamber has an area of high forces while in other FoVs there are very low forces which may render them unusable. Every chip is individual and therefore differs in force and force distribution. This may be due to the coupling to the piezo and due to the geometry of the chip layers which is important for the force generation.

**Figure 3.**
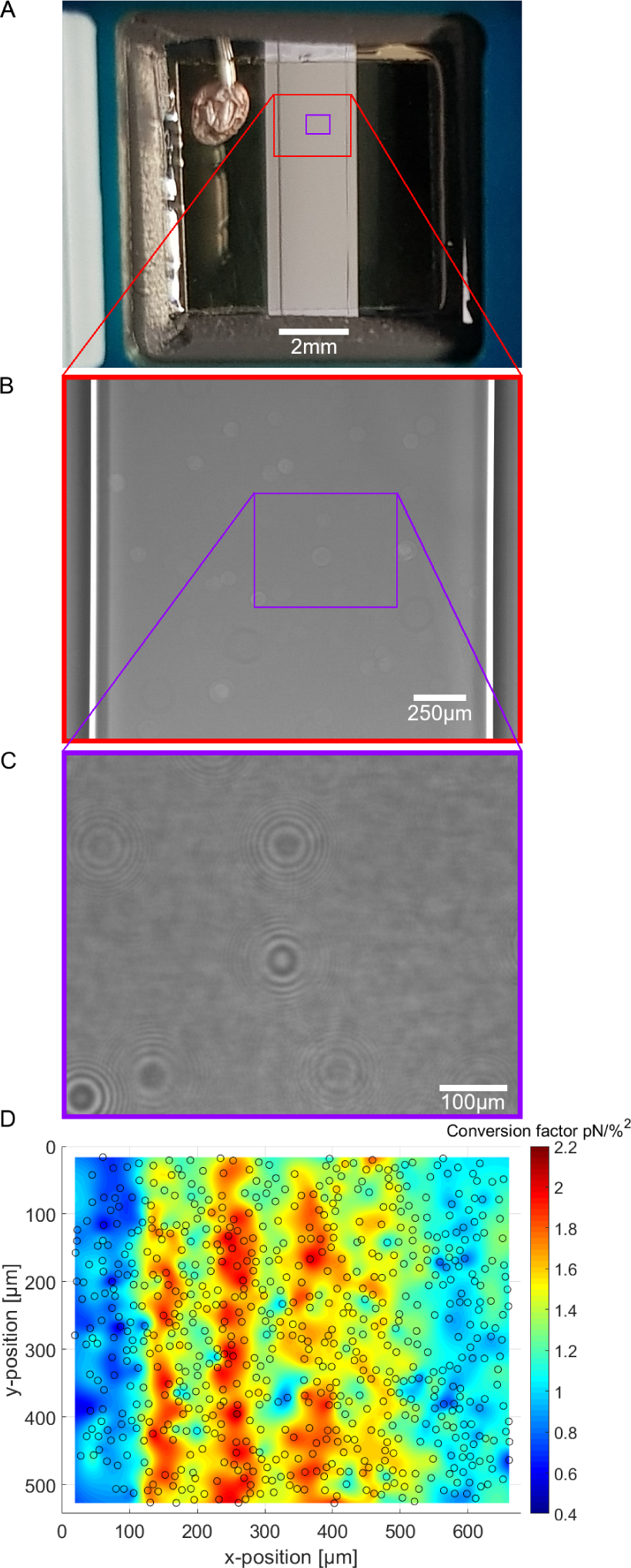
(A) Image of the AFS chip showing the window where the flow chamber and the transparent, quadratic piezoelement can be seen. The connection to the signal generator is placed on the top-left side. (B) Magnified image of the flow chamber showing the flow chamber borders. (C) Further magnified image of the flow chamber showing a typical field of view for the experiments. The random dirt on the piezo is clearly visible. (D) Spatial calibration map (*n* = 1190 beads, merged to 706 beads) obtained by performing the SFC and analyzing with our software *Kitsune*.

### Forces depend on optimal frequency, temperature and medium

From bottom to top, the AFS chip consists of a bottom glass layer, a fluid layer (approximately 100 *μ*m in height), a top glass layer and a transparent piezoelectric element (piezo) glued on top of the glass layer. The glue also serves as a transducer of the acoustic energy to the glass chip. For a fixed chamber geometry design, there is only a given set of usable resonance frequencies. ^42^ The resonance frequency at which the first *z*-node, i.e. the *z*-position where the force equals zero, is between 10 *μ*m to 20 *μ*m is the frequency of the sinusoidal voltage signal applied to the piezo. The performance of the AFS is therefore sensitive to changes in the layer thicknesses. For our chips the acoustic resonance frequency ranges from 14.3 MHz to 14.4 MHz depending on the individual chips. It is expected that the forces decrease if a frequency different to the resonance frequency is used. This is confirmed by performing the SFC to create a spatial map at the same FoV and a fixed temperature for different frequencies in Δ*f* = 0.01 MHz steps (Fig. 4A). As expected, the conversion factor depends sensitively on the applied frequency.

**Figure 4.**
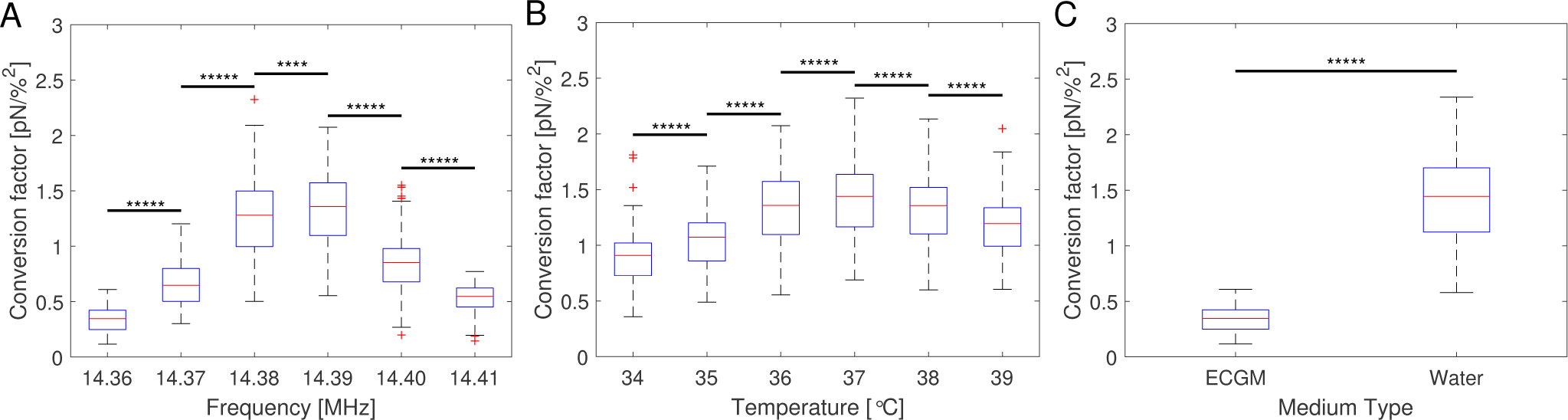
Conversion factors obtained with the SFC with at least 1000 beads (A) for a fixed temperature of 36°C in ECGM at different frequencies, (B) for a fixed frequency of 14.39 MHz in ECGM at different temperatures and (C) for a fixed temperature of 36°C, a fixed frequency of 14.36 MHz in water and in ECGM. All measurements were performed in the same field of view. The significance stars ^****^ represent *p* ≤0.0001, ^*****^ : *p* ≤ 0.00001 using a two-sample *t*-test.

Equation 9 suggests that a change in temperature only results in a change of the absolute value of the acoustic force, as compressibility *κ*_*i*_, density *ρ*_*i*_ of the medium and bead and even the volume of the bead are temperature-dependent. To check this, the SFC is performed at the same FoV and a fixed frequency for different temperatures (Δ*T* = 1°C), while correcting the temperature-dependent viscosity by Eq. A.3. The result shows an optimal temperature (Fig. 4B) for a fixed frequency. This result indicates that for each temperature there is at least one resonance frequency. To test this, the SFC was performed while changing both frequency and temperature. Supplementary Figure S.2 shows that the resonance frequency increases with increasing temperature in the measured temperature range of 25°C - 40°C.

The speed of sound of a material depends on the compressibility and the density of the material (Eq. A.4). This means that the resonance frequency and thus the force will differ in different media. Therefore, for a fixed frequency and fixed temperature the SFC is performed in water and in cell culture medium (endothelial cell growth medium, ECGM) which is used for later experiments. The result clearly shows a difference in the force depending on the medium (Fig. 4C). The median of the conversion factor for water and ECGM appears to vary by a factor of 4.255, when using the same resonance frequency. Supplementary Figure S.3A shows that the resonance frequency inside water is down-shifted compared to the one in ECGM (Fig. 4A). In contrast to previous reports ^41,42^, we find that the acoustic force inside the AFS is dependent on at least seven parameters that may vary in different experiments. It is dependent on the *z*-position shown by the force profile (Fig. 2B), on the *xy*-lateral position indicated by the spatial force map (Fig. 3D), on the applied resonance frequency (Fig. 4A), on the temperature (Fig. 4B) and on the medium (Fig. 4C). Without further testing, the acoustic force will also depend on the type of beads, e.g. polystyrene or silica, as well as their bead sizes inside the range in which Eq. 9 still holds. However, these two bead dependencies may only change the absolute value of the exerted force without shifting the resonance frequency.

### Bead immersion into the cell

To determine the immersion half-angle, 3D stacks of the fluorescently labeled membrane of HUVECs in a monolayer exposed to attached collagen coated polystyrene beads were acquired as described earlier in the materials and methods section. Figure 5A shows the fluorescent signal in the *xy*-plane of highest signal intensity and the *zy*-plane through the bead center for one exemplary bead out of a representative field of view. A 3D-surface plot of the derived cell membrane surface is shown in Fig. 5B. For each bead four different contact points were chosen by eye by estimating the point of turning curvature going outwards from the center of the bead at the deepest indentation. By that, estimating the diameter of the contact circle, the immersion half-angle *θ* was calculated according to Eq. 1. Figure 5C shows the obtained values for every measured bead. The average half-angle was determined to *θ* = 28.73 ± 0.50^0^ (mean ± SEM, n = 43).

**Figure 5.**
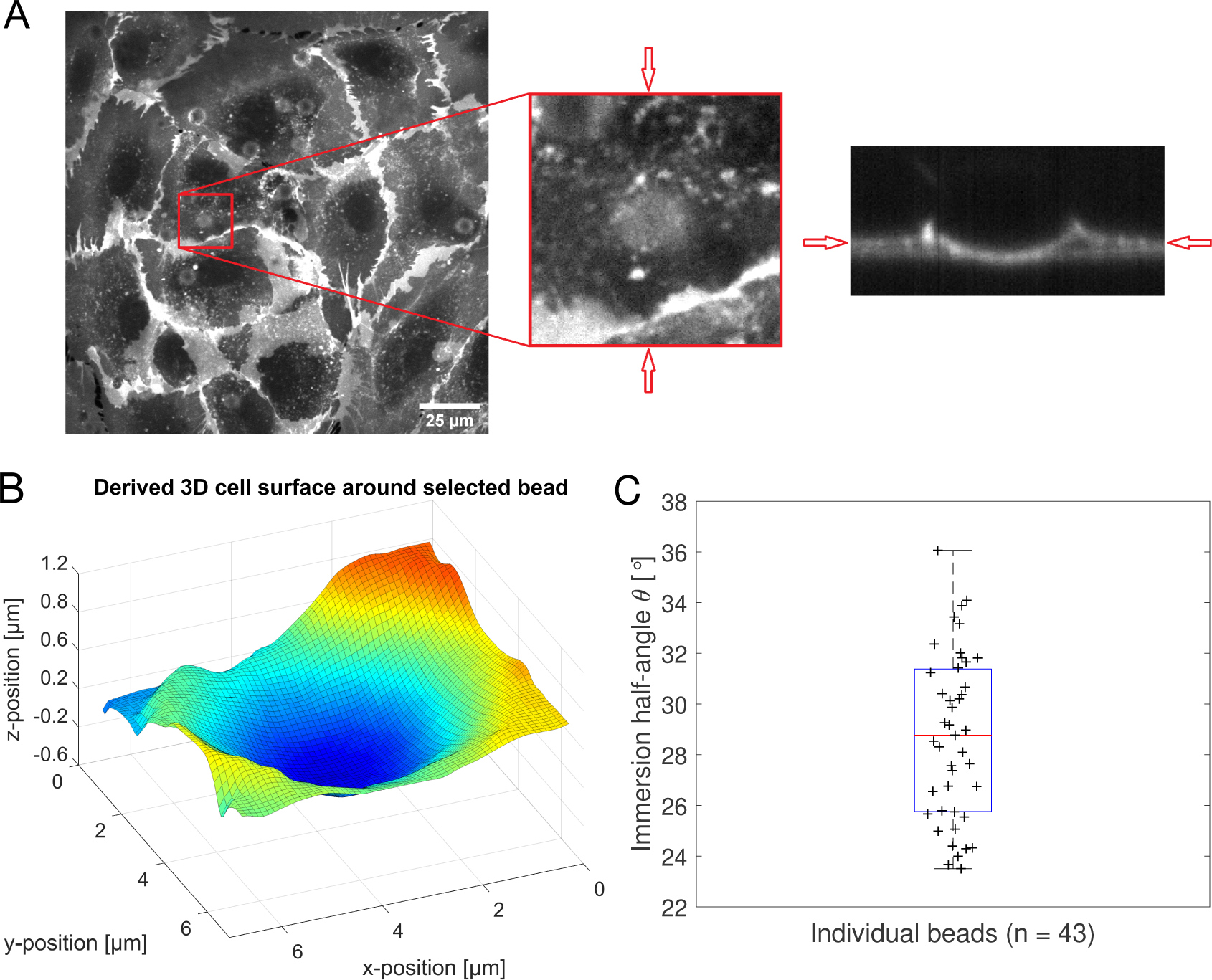
(A): On the left the fluorescence signal of plasma membrane stained HUVECs is shown for the plane of highest intensity of a representative field of view. The middle part highlights a zoomed in selection around an exemplary bead position. On the right the *zy*-slice of the 3D-stack through the bead’s center as indicated by the red arrows is displayed. (B): 3D surface plot of the derived cell membrane surface in close perimeter around the center of the bead from (A), color encodes the position along the Z-axis for better visibility. (C): Distribution of calculated immersion half-angle *θ* of all measured beads.

### Multi-oscillation microrheology using the AFS

To test our new method for microrheology using acoustic forces we oscillate beads attached on a HUVEC cell monolayer. The possibility of culturing a HUVEC monolayer inside the AFS chip with a small volume using our protocol is shown in the Supplementary Information (Fig. S.5). A multi-frequency oscillatory (mOsc) force is applied on the beads with the superimposed frequencies 0.1 Hz, 0.5 Hz and 1.5 Hz. During the experiment the mOsc force remains constant and the medium flow active. The complex shear modulus *G*\*(*ω*) is calculated using Eq. 7. The data was taken from moving time windows of size Δ*t* = 500 s and shift Δ*t*_sh_ = 100 s. The complex shear modulus is calculated at each mOsc frequency.

Fig. 6 summarizes the results of the viscoelastic properties found for the HUVEC cells using the AFS. First, we present the time evolution of the shear modulus’ absolute and phase component of the three different frequencies (Fig. 6A, 6B). As expected for living cells |*G*(*ω*)| increases with frequency, and frequency dependency of the phase hints for a different behavior of the real (*G*′(ω)) and imaginary (*G*″(ω)) part of the modulus regarding the frequency. This is demonstrated in Figure 6C where the temporal mean values of *G*′(*ω*) and *G*″(*ω*) are plotted as function of frequency. The observed crossover suggests that the behavior switches from an elastic to a dissipative regime at a frequency of about 0.2 Hz. This is further demonstrated when plotting the ratio of *G*′/*G*″ (Fig. 6D), that seems to also follow a power-law. An important result shows the log-arithmic plot (Fig. 6C) where two different power-laws can be seen for *G*′(*ω*) and *G*″(*ω*) which would correspond to two different power exponents.

**Figure 6.**
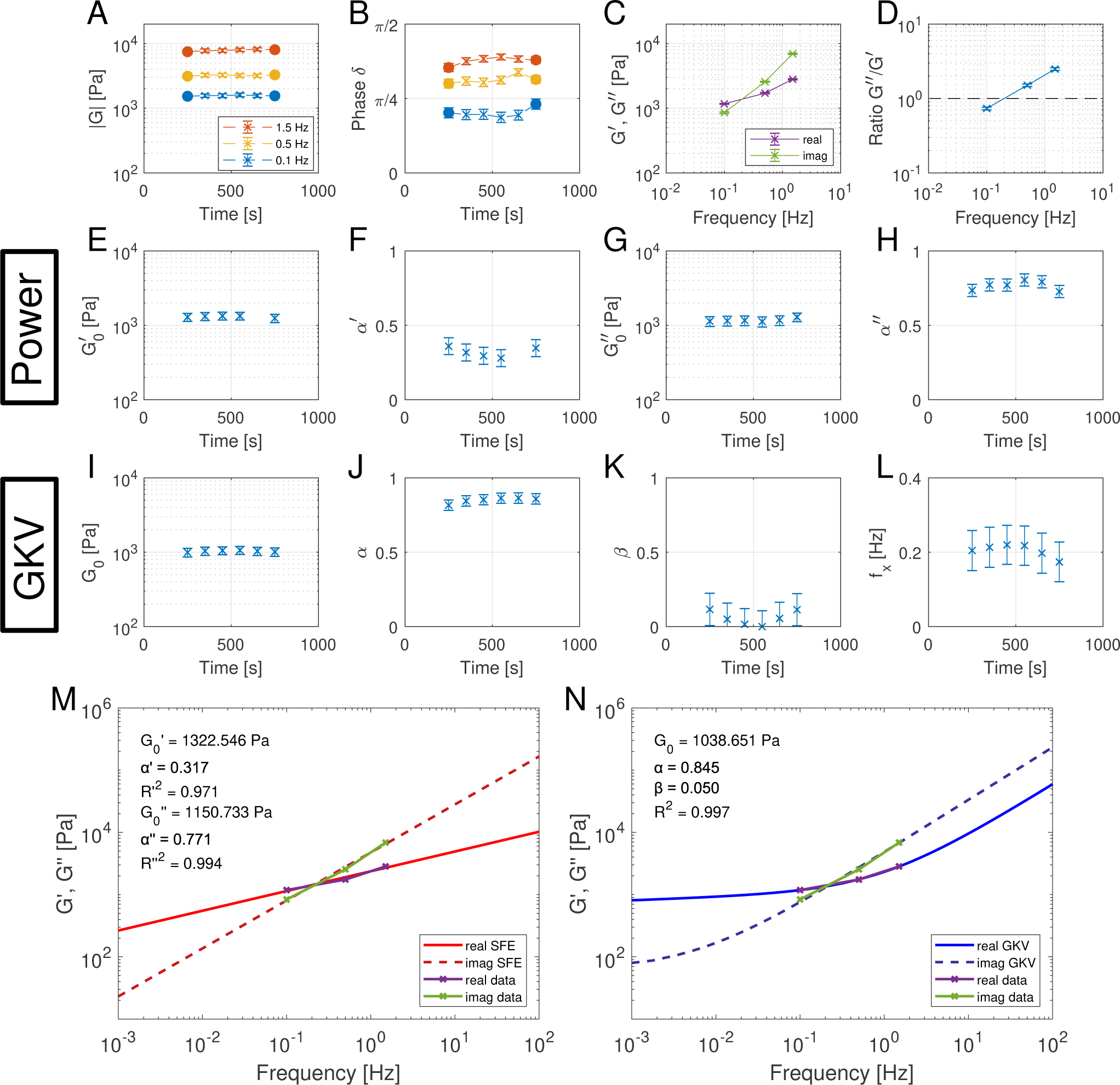
Multi-oscillation microrheology with the SFE model (a power model) and the generalized Kelvin-Voigt (GKV) model. (A, B) Components of the polar form of the complex shear modulus at different time points for three different mOsc frequencies. The filled dots represent independent data results while the c rosses represent data with an overlapping time interval. The evaluation time interval is Δ*t* = 500 s and the shift time is Δ*t*_sh_ = 100 s. The average amount of analyzed beads is *n* = 133 in *N* = 9 experiments. Error bars in (A) represent the standard error of the mean and in (B) from error propagation. (C, D) The mean values of the real and imaginary part and their ratio during the force application for different frequencies. Error bars in (C) represent the standard error of the mean and in (D) from error propagation. (E-H) Fit parameters 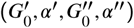 obtained with the SFE model (power model) which is applied separately for the real and imaginary part of the complex shear modulus. Only fit parameters with a goodness of fit o f *R* ^2^ > 0.9 are shown. Error bars represent t he 2 *σ* c onfidence in terval of *n*_b oot_ = 10 0 bo otstrap sa mples. (I-L) Fit parameters (*G*_0_, *α, β*) and the calculated crossover frequency *f*_*x*_ = *ω*_*x*_/2*n* obtained with the GKV model which is applied simultaneously for real and imaginary part of the complex shear modulus. The goodness of fit i s *R* ^2^ > 0.9. E rror bars represent the 2 *σ* confidence in terval of *n*_boot_ = 100 bootstrap samples. (M-N) Representative data of the real and imaginary part of the complex shear modulus at a time point and fit u sing (M) the S FE m odel s eparately for the moduli a nd (N) the G KV model s imultaneously for the moduli.

To compare which power-law model describes the data better we looked at the single fractional element model (SFE) and the generalized Kelvin-Voigt model (GKV). The SFE model can be immediately ruled out as it cannot model crossover events since its expression only consists of one power exponent, meaning the real and imaginary part are parallel in the logarithmic scale and will not cross. To still quantify the slopes seen in Fig. 6C, the SFE is used separately for real and imaginary part and the obtained effectively independent fit parameters for the storage and loss modulus 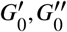 and *α* ′, *α*″ of the SFE are shown in Fig. 6E-6H. These phenomenological power-laws fit the data at all the different time points with a high goodness of fit *R*^2^ ≥0.9. The power exponents of *G*′(*ω*) are clearly different from *G*″(*ω*) while the respective values of *G*_0_ are similar.

To apply a more physically relevant model, we used the GKV model to fit the data simultaneously for both *G*′(*ω*) and *G*″(ω). The fit parameters *G*_0_, *α* and *β* are obtained with Eqs. 4, 5 and are shown in Fig. 6I-6K at different times. The mean fit parameter values are found to be *G*_0_ = 1.033 ± 0.132 kPa, *α* = 0.850 ± 0.035 and *β* = 0.058 ± 0.107 and the resulting mean crossover frequency *f*_*x*_ = *ω*_*x*_/2*n* = 0.204 ± 0.054 Hz. During the mOsc force application the obtained values remain within their error bars which indicate the 2*σ* confidence interval obtained by *n*_boot_ = 100 bootstrap samples. Therefore, we have no indication that the mOsc microrheology changes the cell mechanical properties represented by *G*_0_, *α* and *β* during the measured range. The fit to the data with the independent SFE model and the GKV model is shown in Fig. 6M-6N and with the Generalized Maxwell (GM) model in Supp. Fig. S.8. The GM model is not able to fit the data. Although the GKV model has three fit parameters and therefore a smaller degree of freedom than an independent power-law fit (four fit parameters), it describes the data best and will be used in the following. From the three fit parameters, we can directly deduce the crossover frequency *f*_*x*_ = *ω*_*x*_/2*π* as shown in Fig. 6L.

The results show that mOsc microrheology can be performed with the AFS during a fluid flow as well as without interrupting the mOsc force without affecting the measured cell mechanical properties. Furthermore, the GKV model is fitting the data best.

### Dynamical HUVEC stiffness and power-law depend on the actin cytoskeleton

As we are now able to record time series of microrheology measurements, we turned to a question regarding the dynamical changes of HUVEC viscoelastic properties during manipulation of the actin cytoskeleton. Although it is known that HUVEC stiffness depends on actin, we are now able to perform frequency dependent microrheology measurements and interpret these with the GKV model. We can directly monitor both the mechanical changes, and the timescale of these changes in HUVEC during the disruption of the actin cytoskeleton.

To do so we apply our new method for microrheology with acoustic forces using the mOsc force approach after treating HUVECs with the actin filament polymerization inhibiting drug cytochalasin B (cyto B) which is known to decrease their stiffness at a high concentration, as shown by Stroka and Aranda-Espinoza. ^56^ The experiments are performed as detailed in the Materials and Methods section. Briefly (see Fig. 7), after a 1000 s control measurement we either introduce the treatment (cyto B, red in (Fig. 7C) or a control situation (normal medium, green in Fig. 7A)), where we flush the new medium into the chip by an increased flow for 1000 s. This is followed by a measurement for 1700 s at a low flow rate where we follow the changes of the viscoelastic properties. If the drug has been applied we perform a wash-out experiment by first flushing medium into the chip with increased flow for 1100 s to ensure no drug is left in the chip, and then measure for 700 s. In Fig. 7A-7H the light color represents a low flow rate of 1.66 *μ*l/min and dark colors represent an increased flow rate of 30 *μ*l/min.

**Figure 7.**
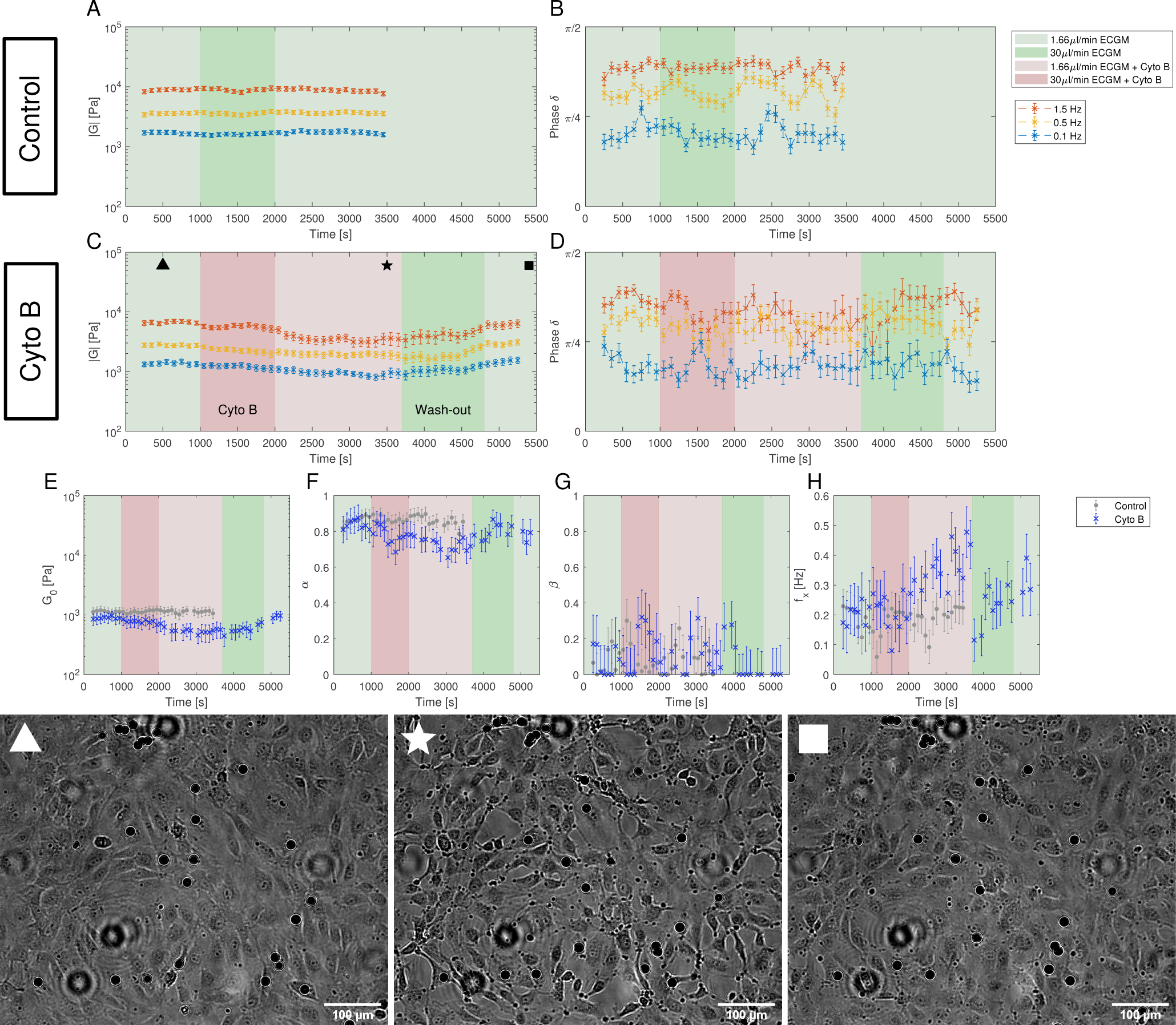
Dynamics of the complex shear modulus measured with multi-oscillation microrheology under flow and cytochalasin B (cyto B) insertion. The marked areas indicate the time intervals of different flow rates with the respective medium composition. All values are shown in Δ*t*_sh_ = 100 s steps at an evaluation time interval of Δ*t* = 500 s. (A, B) The components of the polar form of the complex modulus of a control experiment over 3700 s. For the control experiment only medium (ECGM) has been inserted. The average amount of analyzed beads is *n* = 71 in *N* = 5 experiments. The error bars represent the standard error of the mean. (C, D) The components of the polar form of the complex modulus of the experiment with the addition of cyto B over 5500 s. The cyto B flow has been started at *t* = 1000 s and the procedure to wash it out starts at *t* = 3700 s. The average amount of analyzed beads is *n* = 48 in *N* = 4 experiments. The error bars represent the standard error of the mean. (E, F, G, H) The parameters *G*_0_, *α, β* and *f*_*x*_ = *ω*_*x*_/2*π* obtained with the GKV model for both the control experiment (gray) and the cyto B experiment (blue). The mean fit quality is ^2^ = 0 927, and for all shown fits we have *R*^2^ > 0.7. The error bars represent the 2*σ* confidence interval of *n* = 100 bootstrap *R* .boot samples. (▱) Image of the cells with beads during the measurement at the time point 500 s when no cyto B has been added yet, (*) at the time point 3500 s when cyto B has been inserted and the cells change their morphology and lose cell-cell contacts, (▪) at the time point 5400 s when cyto B has been washed out and the cells regain cell-cell contacts and form a monolayer again. Images are slightly out of focus for the bead tracking and to increase contrast. Additionally, images have been edited to further increase contrast. Scale bar is 100 *μ*m.

The obtained complex shear modulus is shown in Fig. 7A, 7B for a control experiment without addition of cyto B and in Fig. 7C, 7D with the insertion of cyto B. We see a decrease of absolute value in the cyto B case that recovers quickly after removing the drug. The effect of cyto B and the recovery can also be seen by the change of the morphology of the cells (Fig. 7).

Using the GKV model we obtain the parameters *G*_0_, *α, β* and *f*_*x*_ = *ω*_*x*_/2*π* as shown in Fig. 7E-7H. The fit quality is typically above *R*^2^ = 0.9. Supplementary Figure S.10B shows the measurement color-coding the fit quality which shows a decrease of fit quality during the recovery period (4000 s to 4800 s). The parameter *G*_0_ which represents an apparent shear modulus clearly shows a decrease upon adding cyto B as hinted in the values of the complex modulus. The recovery can also be seen by the following increase of *G*_0_ after washing out the drug. The morphology of the cells also reverts back to their initial state. Moreover, the crossover frequency *f*_*x*_ appears to increase under the effect of cyto B (Fig. 7H). Interestingly, the power-law exponent *α* decreases during the treatment, while *β* remains rather unaffected (Fig. 7F, 7G). Therefore, the effect of cyto B may affect the complex modulus at higher frequencies more than in the lower frequencies. In the beginning of the experiment, the control and treatment values remain within the error obtained from bootstrapping.

## Discussion

### AFS as a measurement tool

The first part of the present work provides crucial information for the usage of the Acoustic Force Spectroscopy (AFS) as a measurement tool which is not limited to microrheological experiments. We showed that prior to an experiment the AFS chip has to be calibrated specifically for the experiment in order to gain knowledge of the applied acoustic force. Possibly the most critical information is that the force distribution inside the AFS chip is very inhomogeneous (Fig. 3D). The spatial map of conversion factors shows a force distribution that is highly inhomogeneous but not randomly distributed and thus has to and can be corrected in experiments. Misleading and severely wrong results could occur when comparing experiments at different positions. Therefore, it is very important to do experiments in the same calibrated FoV. The argument of measuring a high amount of beads and then calculating the mean of the result typically would not solve this issue, unless using the same FoV.

Another fact is that there are also lateral forces, see Supp. Fig. S.1. For an attached bead during an experiment, the lateral force is just a low constant force which typically can be neglected. However, for a free bead such as during calibration, this weak lateral force displaces the bead to the lateral nodes. Therefore, the bead tracking should be at a fixed position during the SFC to prevent a calibration of a wrong position. Also, we have shown that the resonance frequency strongly affects the applied forces (Fig. 4A). Although the manufacturer provides a recommended frequency, it should only be regarded as a rough estimate because the resonance frequency depends on many parameters of the experiments, like temperature and medium. For a quick calibration of the resonance frequency at the set temperature and medium, it is sufficient to measure a few beads with the SFC (see Supp. Fig. S.2B) instead of creating several spatial calibration maps as in Fig. 4.

The temperature inside the flow chamber did not change to a high degree (Δ*T <* 0.1°C) during the SFC where the amplitudes are low and the forces are only applied for about 1 s, see Supp. Fig. S.4A. However, at higher amplitudes the temperature will rapidly (minute timescale) rise inside the flow chamber up to a steady state (Supp. Fig. S.4B). As shown, this affects the resonance frequency and thus the forces applied on the bead. Hence, it is important to include knowledge about the temperature during the experiments at high amplitudes. In the current design of the chips the temperature sensor was not located near the flow chamber and thus could not detect changes of the temperature inside the flow chamber. The temperature shown in the LabVIEW software is therefore not the temperature inside the flow chamber, but of the exterior of the chip. During a force application the temperature inside the flow chamber has to be measured with an external sensor prior or during the experiment.

Experiments that require an application of a high amplitude will need to challenge the fact that the forces change during the rise of temperature caused by the applied pressure. This can be solved by applying a long-lasting force so that the temperature reaches a steady state for which the chip has been calibrated. For instance, in our relatively long experiment, the temperature at start of the experiment was set to around 33.5°C. After around 50 s of force application the steady state was reached and the temperature was around 36°C, see Supp. Fig. S.4B. The force at that temperature was then known and did not further change due to the temperature. However, in many other experiments the forces are not applied for a long time. A solution to this could be to apply an acoustic pressure outside of the resonance frequency to exert very low, or hardly any, forces which only increases the temperature by acoustic pressure. Then during the existing acoustic pressure the frequency is changed to the resonance frequency for the given time to apply the desired force. A calibration to determine whether the change in temperature from the one frequency to the resonance frequency is still high should be conducted for this approach. Another solution would be to obtain the temperature-dependency of the force as shown in Fig. 4B. For this approach the change in temperature during the application of the acoustic force has to be known and several spatial calibration maps have to be measured depending on the change in temperature. However, since our experiments were not falling into these categories, they were not further tested.

As previously mentioned, the coupling of the signal to the piezo and the coupling of the piezo to the glass via the glue is important for the applied force. Thus, the performance of the AFS chip can degrade over time which is accelerated inside a humidified incubator. The degradation may result in a weaker force, a change in the resonance frequency or other aspects affecting the applied force. This means that the chip has to be calibrated frequently depending on the surrounding environment.

To summarize the requirements for the usage of the AFS as a measurement tool for experiments, the AFS chips have to be calibrated specifically for an experiment with a set of conditions. A protocol of the SFC is shown in the SI. Typically the choice of medium, bead and temperature are set for the experiment and thus at first, the frequency with a suitable force range has to be found using the SFC. Then, a FoV is chosen which preferably has a high force and a low amount of particles on the piezo that disturbs the tracking. This FoV has to be calibrated by creating the spatial calibration map with the SFC. The temperature change inside the flow chamber by application of the acoustic force has to be measured with an external temperature sensor to ensure that the temperature does not change at a high amount or if it does, then countermeasures as described earlier should be taken. For the analysis of the experiment the lateral position of the bead has to be recorded and compared to the calibration map. The amplitude value sent to the piezo can then be converted to the correct force with the spatial calibration map and possibly a temperature correction.

### A generalized Kelvin Voigt model explains HUVEC viscoelasticity

Using the Acoustic Force Spectroscopy (AFS) for microrheology on a HUVEC monolayer inside the small volume of the AFS chip (6 *μ*l) is possible (see Fig. S.5). However, the long term cell culture inside such a small chip remains challenging because a continuous renewal of the medium is necessary. We find that a very slow flow of 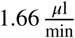, corresponding to a wall shear stress of 5.9 mPa is sufficient. Without continuous medium exchange, the cells do neither spread nor form a monolayer (see Supp. Fig. S.6).

In order to perform the multi-oscillation (mOsc) microrheology beads were inserted and attached to the HUVECs. Only beads that were strongly attached to the cells and did not detach due to flow shear stress nor application of the force were tracked. The effect of different bead coatings might be of interest for further measurements.

HUVEC cell monolayers are optimal cells for the AFS measurements regarding their flat spreading which leads to a low cell contribution in the bright-field images, hence enabling excellent bead tracking on their surface. Additionally, the height of HUVECs usually does not exceed 3 *μ*m at the cell body and is 1 ≥ *μ*m at cell periphery ^57^, and thus, the force profile in *z* obtained from the calibration can be regarded as constant inirection (Fig. 2B). However, due to the motility of the HUVECs the lateral position of the beads also changed during the measurement. This needs to be accounted for by correction of the calibration factor during the measurement using the calibrated spatial conversion factor.

In the frequency dependent mOsc microrheology measurements we span more than one order of magnitude in a single measurement (0.1 Hz, 0.5 Hz and 1.5 Hz). Using more frequencies is possible but either leads to lower signal per frequency or requires an increase in amplitude which also increases the local heating. Higher frequencies would require a faster camera for tracking. Up to now, no frequency dependent microrheology experiments have been performed on HUVEC monolayers. We find a power-law rheology consistent with previous measurements on cells ^23,26^, and the overall elasticity values in the kPa regime comparable to previous measurements using AFM. ^22,58^ Favorably, the choice of the frequency values enabled the resolution of a crossover event at which the real and imaginary part of the complex shear modulus are equal (Fig. 6C). Classical models using one single fractional element to model power-law materials fail to explain such crossovers. On the contrary, the generalized Kelvin-Voigt (GKV) model, comprised of two single fractional elements in parallel, was able to provide an excellent fit to the data. Here, the full complex curves are explained by only 3 parameters. The found values of *G*_0_ can be interpreted as steady state elasticity, as the second power-law exponent *β* is found to be close to 0 and in the long time limit the elastic modulus becomes independent of the frequency (see Supp. Fig. S.8). The found elastic shear modulus of *G*_0_ = 1.03 ± 0.13 kPa corresponds to a Young’s modulus of *E* = 3.09 ± 0.39 kPa assuming a Poisson ratio of 0.5. This measurement is in excellent agreement with previous measurements of HUVEC cortical stiffness at cell body using AFM yielding a Young’s modulus in the range of 2.70 − 3.23 kPa ^59–62^, but is higher when compared to another study that found *E* = 0.757 ± 0.016 kPa at cell body and *E* = 1.042 ± 0.014 kPa at cell periphery. ^56^ Thanks to the frequency dependent measurements we determine a crossover frequency which suggests that the mechanical properties at the membrane of HUVECs undergo a solid to liquid transition in the second timescales. This is at a much slower time scale as reported for other cell types where the transition on the cortical level happens in the millisecond regime. ^63^ Whether this is relevant for the function of endothelial cells that are exposed to variable shear stress has to be determined in further studies. Overall, we showed that the mOsc microrheology can be applied using acoustic forces and that the data can be analyzed with the GKV model which yields an apparent shear modulus *G*_0_ and the crossover frequency *ω*_*x*_ obtained by the two power exponents (*α, β*). Our results are in excellent agreement with literature and provide new insights in the viscoelastic properties of HUVEC monolayers.

The advantage of the AFS is that it makes use of a microfluidic chip, meaning a closed system that allows a fluid flow. Drugs, such as cytochalasin B (cyto B), can be flushed inside during the measurement, i.e. during the application of the force, to capture the dynamics of the complex shear modulus without stopping the measurement. Although the effect of disruption of the actin cytoskeleton on cell mechanics is well established, direct monitoring during the application, and measurements of HUVEC mechanics induced by fluid flow induced shear forces have not been possible previously. Our results show that long (more than 3700s) application of the mOsc force did not affect HUVEC mechanics. It is reported that shear flow has an effect on cell mechanics at a high shear stress, e.g. 2 Pa ^64^, however, in our switch from low to high fluid flow for the treatment insertion, the wall shear stress only increased from 5.9 mPa to 106.1 mPa. As it is shown that wall shear stress of 320 mPa ^65^ for 24 h did not induce cell alignment or change in protein expression we expect no effect due to the induced forces. In our experiment, we also did not observe changes in the viscoelastic properties during the recording time (see Figure 7A-7B). In contrast, upon application of cyto B, the complex shear modulus decreased significantly (Fig. 7C-7D) which also could be captured using the GKV model shown in the significant decrease of the apparent shear modulus *G*_0_ to almost 50% of its initial value (Fig. 7E and Supp. Fig. S.10A).

These experiments are done in ECGM, and the calibration of the FoV done in water has been corrected using the scaling factor for water to ECGM obtained from Figure 4C. This scaling factor was obtained by the ratio of the median values of the calibration maps. The mOsc microrheological experiment required beads that did not detach and could be tracked for a long time. Therefore, unlike the short SFC measurements the amount of beads per experiment is limited (*n* ≈ 15). Unfortunately, this means that the usual high-throughput advantage of the AFS does not apply for the present microrheological experiment in our hands. However, considering long experiments with changes in the conditions that affects the sample globally, such as the insertion of a drug, the AFS is highly suitable in regards of measuring multiple beads simultaneously.

## Conclusions

Overall, we establish a novel method for multi-oscillation microrheology on cell monolayers using acoustic forces with the Acoustic Force Spectroscopy (AFS) which alled us to deter-mine the frequency dependent viscoelastic properties of HUVEC monolayers. The data is well explained with a generalized Kelvin Voigt model using fractional elements. However, the usability of the AFS technique for such measurements is still rather limited in the current situation. Measurements with the AFS require a thorough calibration depending on the experiment, mainly due to the force inhomogeneity and the dependencies of the applied force on the applied frequency, temperature and medium. This result of our work is crucial for most current and future experiments with the current design of the AFS. While the advantage of high-throughput measurements does not apply for the present microrheological experiment, the advantage of the flow system enabled the first measurement of the dynamics of the complex shear modulus of HUVEC monolayers under flow, and to determine the time evolution of a drug effect on these cells. Hence, we have opened the door for more complex experiments of microrheology under flow that are typically not possible with other techniques like AFM. Future experiments could be to measure the dynamics at different flow stresses, and the time course of such mechanical stresses, as the mechanical response to shear stress of endothelial monolayers is a key element relevant for vascular integrity and immune response.

## Appendix

### Temperature-dependent properties of water

The temperature-dependent density of air-saturated water *ρ*_AS_(*T*) and isothermal compressibility *κ*(*T*) can be approximated with a polynomial of fourth order ^66^

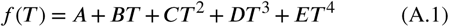

with the constants shown in Table A.1 and the temperature *T* ∈ [5, 40]°C. The compressibility-corrected temperature-dependent water density is then given by ^66^

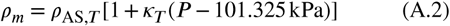

with the ambient pressure *P* in kPa. The viscosity is also temperature-dependent, *η* = *η*(*T*), and for water it can be approximated with a modified Andrade equation given by ^53^

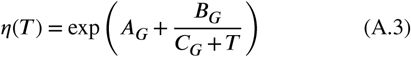

with the constants *α*_*G*_ = −3.63148, *B*_*G*_ = 542.05 °C, *C*_*G*_ = 129.0 °C and the temperature *T* in [°C]. The speed of sound in water can be calculated with the Newton-Laplace equation

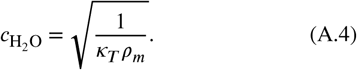

### Viscosity correction due to the surface boundary

The viscosity correction by Faxén’s law for translation parallel to the boundary surface is given by ^21^

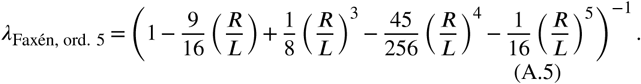

An approximation of the viscosity correction by Brenner (Eq. 13) for translation perpendicular to the boundary surface is given by ^20^

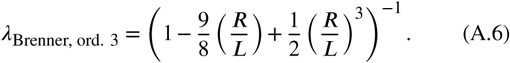

A higher order approximation of the viscosity correction by Brenner may yield a better result.

## Acknowledgements

We thank Felix Oswald from LUMICKS for useful discussions about the AFS. We gratefully acknowledge the support of the Cells in Motion Cluster of Excellence (EXC 1003 CiM) and the financial support of the European Research Council (ERC-CoG, PolarizeMe 771201). We also thank the Interdisziplinäres Zentrum für Klinische Forschung (IZKF, Bet1/013/17) of Münster and Prof. Dr. V. Gerke (Institute of Medical Biochemistry, University of Münster, Münster, Germany) for the HUVEC culture.

---

**Table A1.** Values of the constants to calculate the temperature-dependent density and isothermal compressibility ^66^.

| Property | A | B $\left[\frac{1}{^\circ\text{C}}\right]$ | C $\left[\frac{1}{^\circ\text{C}^2}\right]$ | D $\left[\frac{1}{^\circ\text{C}^3}\right]$ | E $\left[\frac{1}{^\circ\text{C}^4}\right]$ |
| --- | --- | --- | --- | --- | --- |
| Density $\left[\frac{\text{kg}}{\text{m}^3}\right]$ | 999.84847 | $6.337563 \cdot 10^{-2}$ | $-8.523829 \cdot 10^{-3}$ | $6.943248 \cdot 10^{-5}$ | $-3.821216 \cdot 10^{-7}$ |
| Compressibility $\left[\frac{1}{\text{kPa}}\right]$ | $50.83101 \cdot 10^{-8}$ | $-3.68293 \cdot 10^{-9}$ | $7.263725 \cdot 10^{-11}$ | $-6.597702 \cdot 10^{-13}$ | $2.87767 \cdot 10^{-15}$ |

## Supplementary Information

### Stokes Force Calibration (SFC)

#### Theoretical aspects of acoustic radiation force on small particles

The acoustic radiation force *F*_rad_ on a small spherical particle in a standing acoustic wave in a viscous fluid is the negative gradient of the potential *U*_rad_ and is given by ^51^

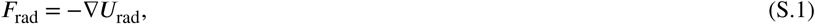

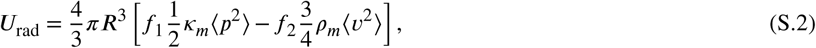

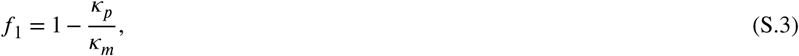

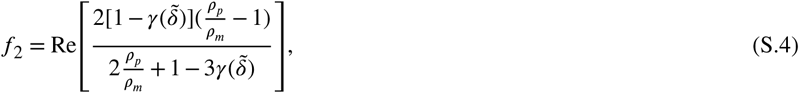

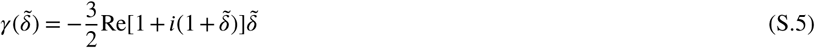

with the particle radius ***R***, compressibility *κ*_*p*_ and *κ*_*m*_, density *ρ*_*p*_ and *ρ*_*m*_ of the particle and surrounding medium, respectively. The time average is denoted by ⟨·⟩. with the acoustessure *p* and the acoustic velocity *v*. The dimensionless parameter 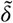 is given by 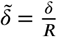 with the viscous penetration depth 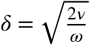 and the kinematic viscosity 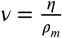 given by the ratio of the dynamic viscosity *η* and the density of the medium, and the angular frequency *ω*. Similar to the view in ^51^, for the case in this work the viscous penetration depth is approximately *δ* = 0.15 *μ*m for a 14.3 MHz sound wave in water at room temperature. The particle radius used in this work is 5 *μ*m and thus ^51^ sufficiently large to be described by an inviscid theory where

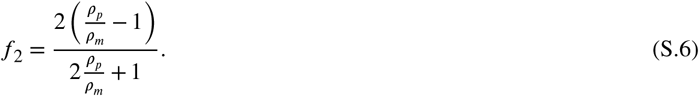

For a one-dimensional planar standing wave the incoming acoustic pressure and velocity field are given by

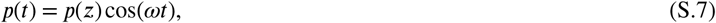

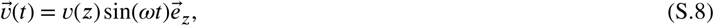

respectively, with the pressure *p*(*z*) and velocity *v*(*z*) at the position *z*. With Eq. S.1 and the time averages 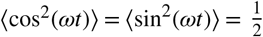 the resulting one-dimensional acoustic radiation force in *z* is then ^42,52^

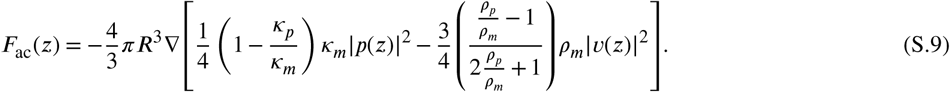

#### Lateral translation of free beads

A lateral translation of free beads can be observed when applying an acoustic pressure using the AFS. This can be seen in Supp. Fig. S.1 and in Movie 1.

**Figure S.1.**
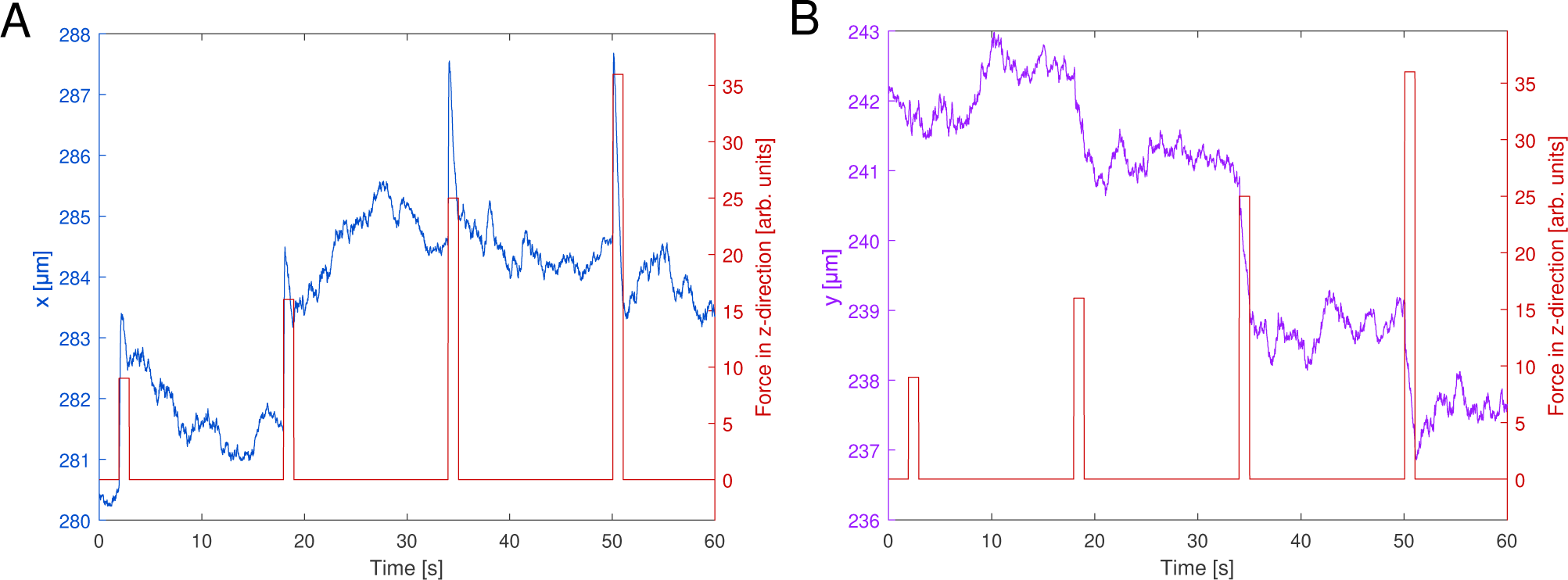
The lateral-positional data of the beads during the Stokes Force Calibration. During force application the beads show a lateral translation. The absolute values of the positional data refer to the position of the bead inside the field of view.

#### Data analysis of the SFC

In order to obtain a correct calibration map some beads were filtered out using our software *Kitsune* (available on request); the filter criteria are presented in the following. The raw data obtained with the modified LabVIEW software is an ASCII-file containing the *xyz*-position, the applied amplitude value *V*_%_ and the frequency *f* at a time point *t* and a force type value *F*_*t*_ with *F*_*t*_ = 0 at time points when no voltage signals were sent to the piezo and *F*_*t*_ = 7 for time points when a pulling force calibration and *F*_*t*_ = 5 for a multi-oscillation force. For the SFC analysis the time points with *F*_*t*_ = 7 were analyzed. The first time point with *F*_*t*_ ≠ 0 was set to *t* = 0 s. The *z*-data was shifted by an offset such that *z*(*t* = 0) = 0 *μ*m. Firstly, the following criteria had to be fulfilled during the applied constant force before being further analyzed.

- There are no *z*-data points that equal zero which typically indicate tracking errors.
- There are no sudden changes in *z*, i.e. between each sampling point the difference must be lesser than 10 *μ*m.
- The maximal *z*-data must differ by at least 5 *μ*m above the surface to filter out attached beads or weak forces that could not lift the bead as high during the force application time of 1 s.

The relevant *z*-positional data is from the ground to the *z*-node, i.e. the *z*-position at which the force equals zero. However, the acoustic forces inside the AFS is typically also exerted laterally. Therefore, the bead is also displaced laterally. The force is typically still applied even when the bead has reached the *z*-node, but due to a possible slight tilt of the flow chamber or a *xy*-position-dependent *z*-node the *z*-position of the bead may gradually change. This is taken into account by only analyzing the start-point up to the time point when the bead first reaches the *z*-node. For the estimation when the *z*-node is reached the following steps were performed.

- The full uncropped range is used and Eq. 16 is used to obtain the optimal fit parameters (see description in the Results section).
- The obtained velocity is numerically differentiated to obtain the acceleration.
- The first *z*-node estimation is the *z*-position at the time point when the absolute of acceleration is minimal.
- If the first *z*-node estimation is *<* 13 *μ*m it is set to 13 *μ*m.
- New optimal fit parameters are now obtained with the *z*-data from the start point to the first *z*-node estimation.
- If the fit quality, here, represented by the sum of the squared difference of the numerically obtained *z*_num_ and the measured *z*_meas_, is greater than 1 *μ*m^2^ the *z*-node estimation is decreased by 0.1 *μ*m until the fit quality is smaller than 1 *μ*m^2^ or the *z*-node estimation is smaller than 13 *μ*m.

With the obtained fit parameters the acoustic force profile can be calculated with Eq. 9. The calibration values are saved to be run through the following filters to create the final calibration map.

The values obtained from that amplitude for that bead are filtered out if at least one of the following criteria of *Filter-Type 1* is fulfilled:

- The fit quality (represented by the sum of the squared difference of the numerically obtained *z*_num_ and the measured *z*_meas_) is greater than 2 *μ*m^2^.
- The fit quality is smaller than 0.001 *μ*m^2^.
- The *x*-position or the *y*-position of the bead is *<* 0 *μ*m.
- The force at 1 *μ*m changes more than 15 % of the maximal force.
- The force at 1 *μ*m is lesser than 0 pN.
- The force at 1 *μ*m and the maximal force is greater than 80 pN.

After running through the *Filter-Type 1* the next filter is *Filter-Type 2*. Typically, there are at least three different amplitude values for each bead. *Filter-Type 2* filters out amplitude values that yielded a lower force, albeit being a higher value than the previous amplitude value for one bead. The next final *Filter-Type 3* removes values of all beads that only have one amplitude measured after all the previous filters to ensure the correct quadratic fit for the conversion factor where at least two data points are needed. For the remaining calibration values the conversion factors are obtained for each bead by fitting Eq. 17.

Due to the high amount of beads that were not simultaneously tracked they might overlap. Therefore, the mean values of the conversion factor of the beads that are closer than a merge distance of 10 *μ*m are used. The map is then created by creating a rectangular grid with mesh size of 0.5 *μ*m starting from the minimal measured *xy*-position to the maximal measured *xy*-position. The grid interpolation is based on a biharmonic spline and is performed by the in-built MATLAB function *griddata*.

#### Temperature-dependency of the resonance frequency

A short version of the SFC was performed inside a field of view with more than 14 beads at different locations. Only one amplitude was used, therefore the force is represented instead of the conversion factor. The measured data were in Δ*T* = 1°C temperature steps and Δ*f* = 0.01 MHz frequency steps. The mean force of more than 14 beads at different positions in one field of view is shown in Supp. Fig. S.2 with a cubic interpolation using the in-built MATLAB function *griddata*. Supplementary Figure S.2 shows that the resonance frequency depends on the temperature. Moreover, the resonance frequency increases with higher temperatures.

**Figure S.2.**
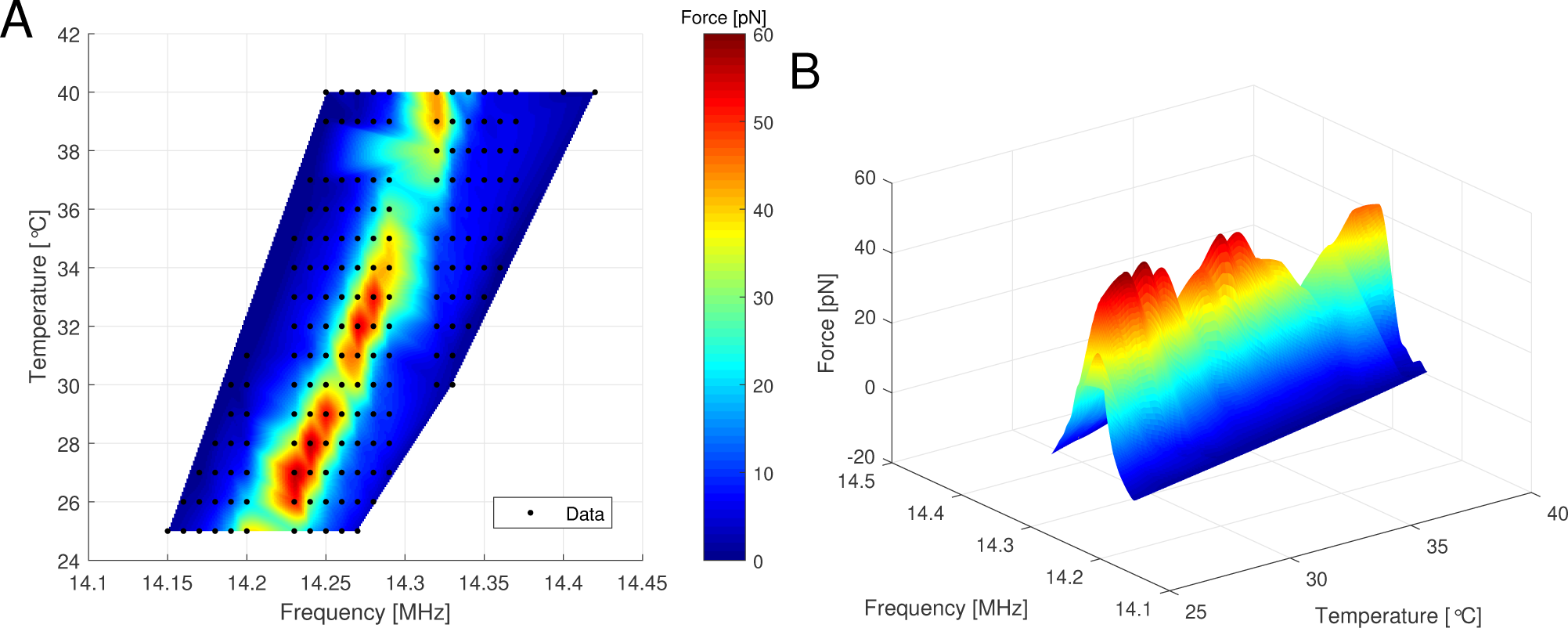
Interpolated mean forces on more than 14 beads at different temperatures and different frequencies. (A) 2D-view of the relationship between frequency and temperature. The resonance frequency increases with higher temperature. The black dots present measured data. (B) 3D-view showing the forces at a frequency and temperature.

#### Resonance frequency shift inside different medium solutions

At a fixed frequency and temperature, the force depends on the medium as shown in Fig. 4C. For a fixed temperature of *T* = 36°C the resonance frequency in ECGM was about *f*_res, ECGM_ = 14.39 MHz for our calibration chip. Supplementary Figure S.3A shows that the resonance frequency inside water is about *f*_res, water_ = 14.35 MHz and thus down-shifted compared to the one in ECGM. There is still a difference of the conversion factors in ECGM and in water at a fixed temperature of *T* = 36°C at their respective resonance frequencies, however, it is not too high (Supp. Fig. S.3B) and may be due to the resolution of the exact resonance frequency scan, as it was performed at Δ*f* = 0.01 MHz steps.

**Figure S.3.**
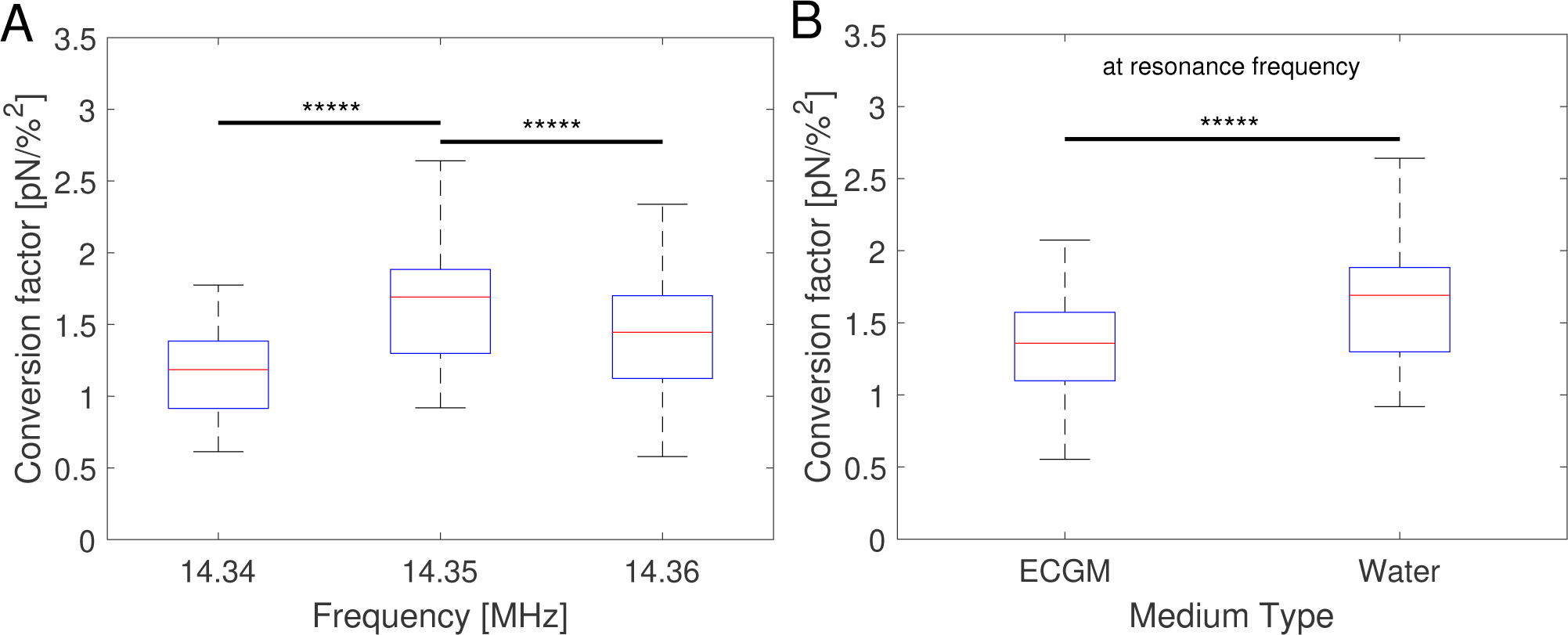
Conversion factors obtained with the SFC with at least 1000 beads (A) for a fixed temperature of 36°C in water at different frequencies, (B) for a fixed temperature of 36°C in ECGM at the resonance frequency *f* = 14.39 MHz and in water at the resonance frequency *f* = 14.35 MHz.

#### Temperature change during force application

It is expected that the temperature changes upon force application. The in-built temperature controller inside the chip holder cannot measure the temperature inside the fluid chamber, but measures the chip’s exterior instead. Therefore, an external temperature sensor (disassembled from an AFS G1 chip holder) is used to measure the temperature of the bottom glass underneath the fluid chamber. For this, thermal grease is spread onto the bottom glass. However, care has to be taken of that the thermal grease or the sensor may leave scratches at the bottom of the chip which renders the affected FoVs useless. In addition, if thermal grease is used, the interface glass to air is then changed to glass-grease which typically increases the acoustic energy transmission and thus results in a weaker force. This can lead to a lower temperature increase compared to the situation without the thermal grease.

During the SFC the temperature hardly changes (Supp. Fig. S.4A). The exact time of the force application during the measurement of 90 s was not known, however, no relevant temperature change could be measured.

When applying higher amplitudes, such as during the mOsc experiment, the temperature drastically increases (Supp. Fig. S.4B). From around 33.5°C the mOsc force increased the temperature to a steady state temperature of around 36°C after about 50 s. Due to the oscillation, the temperature also changed by about Δ *T* = 0 2°C which is negligible. The high flow rate of 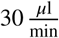 of medium at room temperature only slightly decreased the temperature by Δ*T <* 0.5°C which is also negligible.

**Figure S.4.**
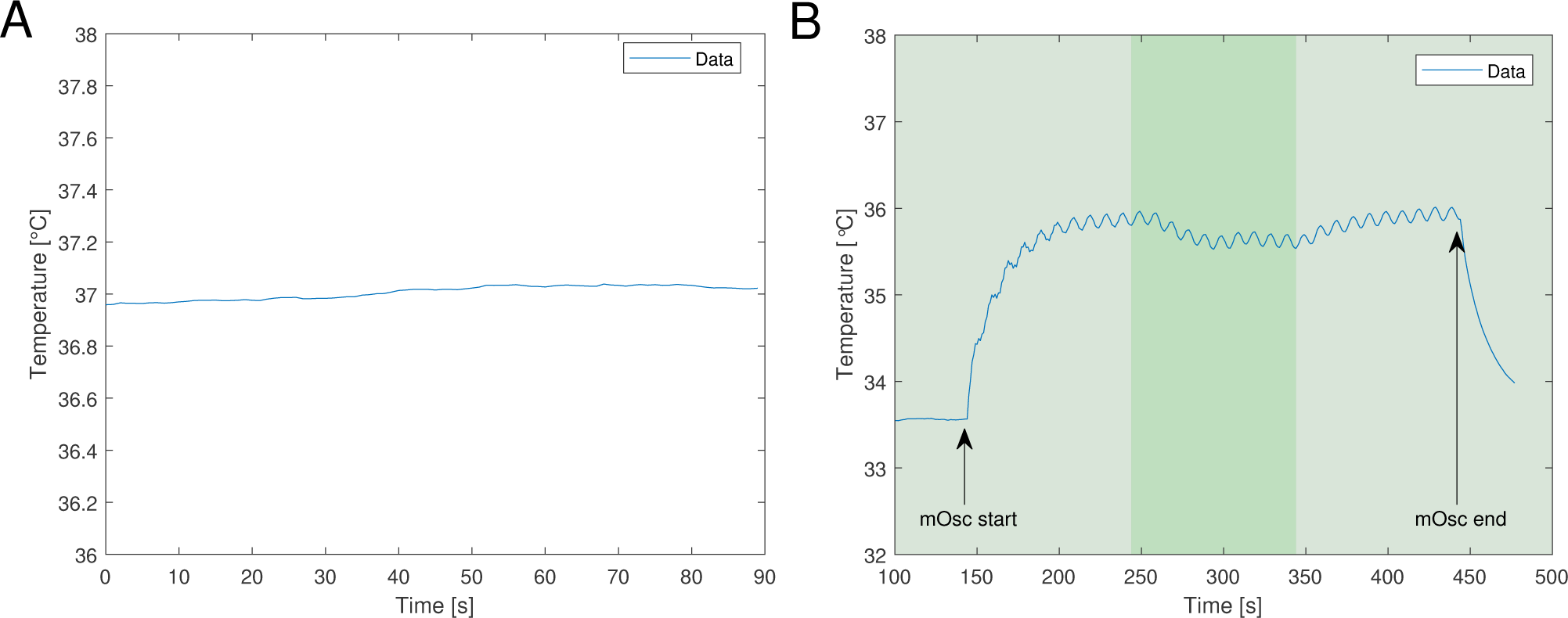
Temperature change during force application and flow measured with an external temperature sensor. (A) The temperature during the SFC. The exact time of the SFC forces were unknown, however, inside the force application window (measurement time of 90 s) there is no visible temperature change. (B) The temperature during the mOsc force application. The light green area represents the slow flow of medium 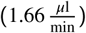 and the flow rate in the green area is 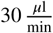. The mOsc force causes an increase in temperature up to a steady state temperature with a negligible oscillation. The higher flow rate s lightly d ecreased the temperature.

#### HUVEC culture inside an AFS chip

To demonstrate the possibility of culturing a HUVEC monolayer inside the AFS chip, the cell development is recorded about three minutes after seeding with a sampling time of 1 min. For the recording, the chip is placed on the AFS equipment with the temperature controller instead of the incubator. Due to the closed flow system, the CO_2_ environment is not of importance. Fig. S.5 shows selected time frames and Movie 2 shows the culture inside the AFS every minute. Inside the AFS chip HUVECs started to spread out onto the surface after about 2 h and were fully spread out a day after seeding. In this case, after 93 h a monolayer of HUVEC had formed. During the incubation, the medium has been renewed with a flow rate of 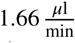. The medium renewal was essential to culture HUVEC monolayers inside the AFS (see Supp. Fig. S.6 and Movie 3). Thus, we show that a monolayer of HUVECs can be cultured inside an AFS chip using our protocol.

**Figure S.5.**
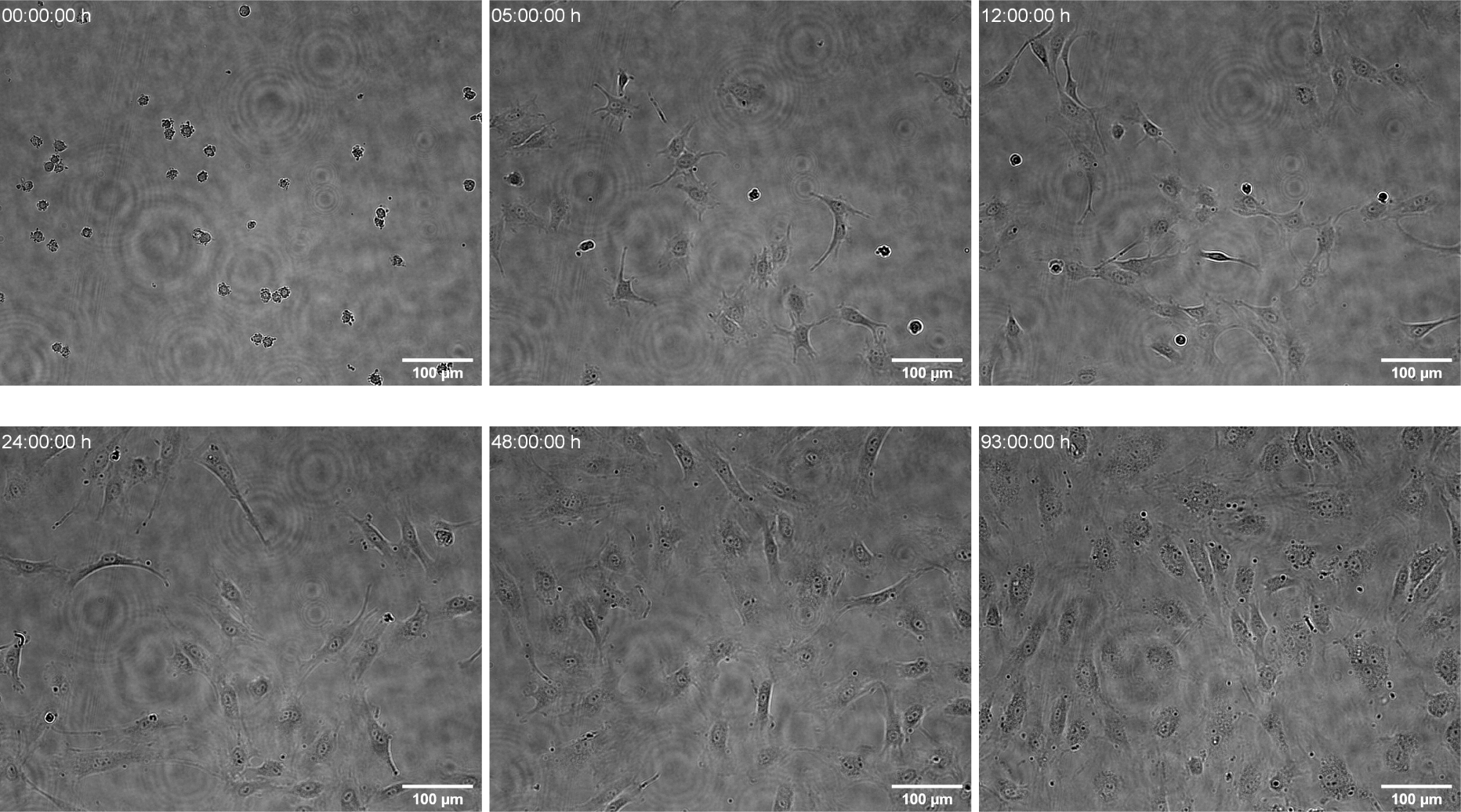
HUVEC culture development inside the AFS chip at different time frames. The bright-field images are slightly out of focus to increase the contrast of the cells. Cells are still round three minutes after seeding (*t* = 0 h). Cells are fully spread out onto the surface and proliferated (*t* = 48 h). A monolayer has formed (*t* = 93 h). The images were edited to further increase contrast. Scale bar is 100 *μ*m.

Without the renewal of culture medium the cells die before being able to form a monolayer (Figure S.6, Movie 3).

**Figure S.6.**
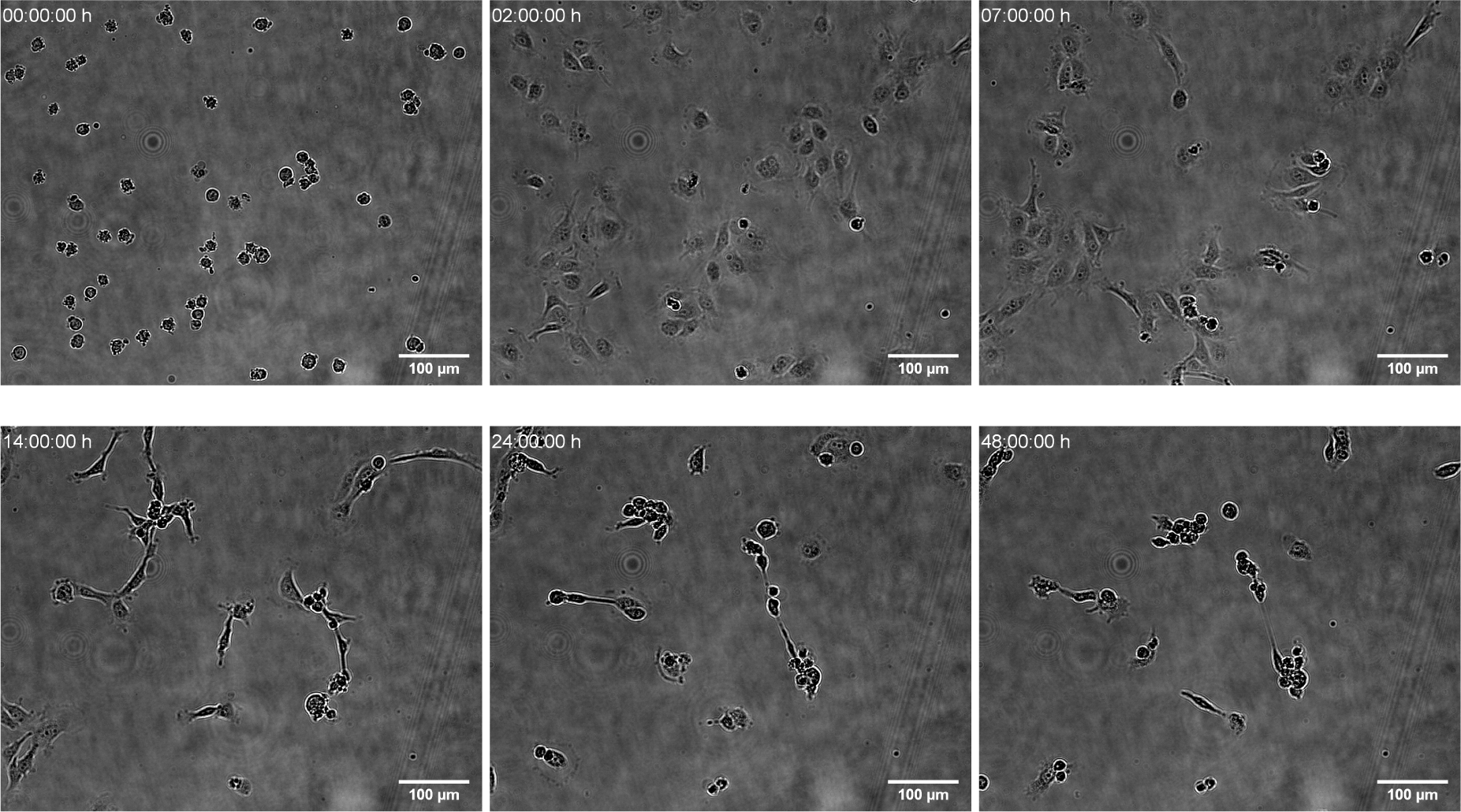
HUVEC culture development inside the AFS chip at different time frames without medium renewal. The bright-field images are slightly out of focus to increase the contrast of the cells. Cells are still round three minutes after seeding (*t* = 0 h). After *t* = 2 h the cells are spread out onto the surface. After *t* = 7 h the cells start to change their morphology and become elongated. At *t* = 14 h the elongated cells can be clearly seen. After *t* = 24 h the cells round up and die. After *t* = 48 h most of the cells died and the cells are unable to form a monolayer. The images were edited to further increase contrast. Scale bar is 100 *μ*m.

### Microrheological Experiment

#### Theory of microrheology

The mechanical properties of complex materials can be measured using microrheological experiments. To interpret the obtained data the microrheological analysis undergoes a few assumptions as follows. The first assumption is that the cell is behaving as a linear viscoelastic material. This can be considered valid for a sufficiently small deformation of the material. For a linear material the superposition principle holds, thus the relation of the strain *ϵ*(*t*) and the applied stress *σ*(*t*) has the form of the hereditary integral

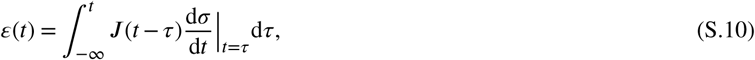

with the creep compliance ***J*** (*t*). The Laplace-transformation of Eq. S.10 yields

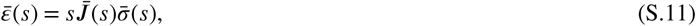

with the complex frequency *s* in the Laplace-domain, or in the Fourier-domain identify *s* = *iω*, hence

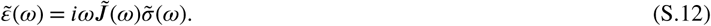

Considering an oscillatory applied stress 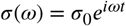 of frequency *ω* = 2*πf* the resulting steady state strain is also oscillatory 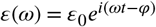 with the same frequency *ω*, but with an additional phase shift *ϕ*. For a purely linear elastic material represented by a single spring element, the phase shift equals zero (*ϕ* = 0°) and the strain immediately follows the applied stress; for a purely linear viscous material, a single dash-pot element, the strain lags behind the stress by *ϕ* = 90°. Here, the material of interest is viscoelastic and the phase shift is therefore *ϕ* ∈ (0°, 90°). The viscoelastic complex modulus *G*\* can be defined as the ratio of the stress to strain

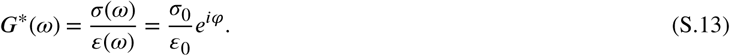

The relation between the viscoelastic complex modulus and the creep compliance with Eq. S.12 is then

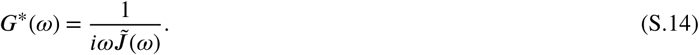

The second assumption is the choice of the model that describes the viscoelastic material. In this case, a model based on fractional calculus is chosen. The following calculations are mostly based on ^47^. The underlying assumed element is that the creep compliance increases as

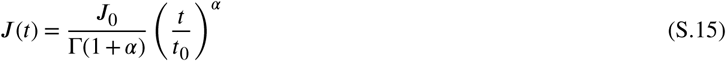

with an arbitrary time constant *t*_0_, a constant ***J***_0_ and the complete Gamma-function Γ (·) for *α* ∈ [0, 1]. With Eq. S.10 and Eq. S.15 the strain is given by

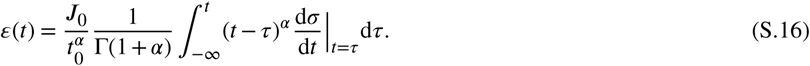

With *α* = *γ* −1 identify the special form of the Riemann-Liouville integral, also known as the Weyl’s fractional integral 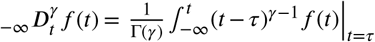, so that

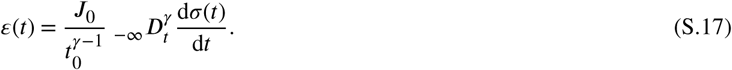

The fractional integration and differentiation can be obtained, shown by performing fractional integration followed by ordinary integer differentiation, see Schiessel *et al*. ^47^, and Weyl’s fractional integral can be notated as 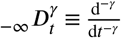. Re-substituting *γ* = *α* +1 yields the rheological constitutive equation (RCE)

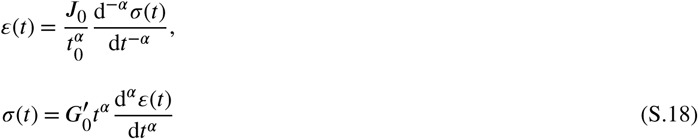

using the fundamental relations 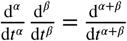 and 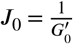. As coined by Schiessel et al.^47^ the RCE represents the single fractional element (SFE). The complex modulus can be obtained by Fourier transforming Eq. S.18

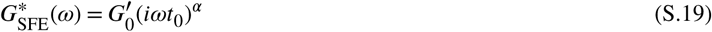

and with 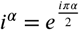 and comparing with Eq. S.13 the previously described phase shift is

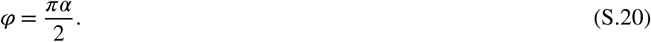

Thus, for a purely linear elastic material the exponent *α* equals zero, for a purely viscous material, *α* = 1, and for a viscoelastic material *α* ∈ (0, 1). Due to the scaling invariance of Eq. S.19 the time constant is set to *t*_0_ = 1 s and 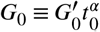. Note that the RCE and the resulting complex shear modulus for the SFE (Eqs. S.18, S.19) represent a power-law behavior as often ^37,48^ used to describe linear viscoelastic materials. The complex modulus can be represented in real and imaginary parts 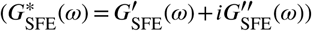 or in polar form with the absolute and phase 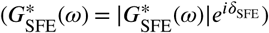:

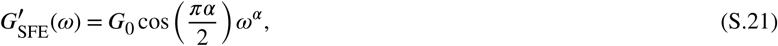

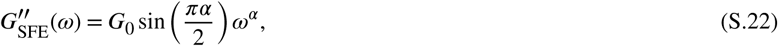

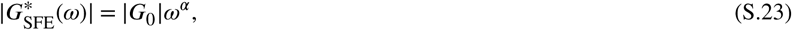

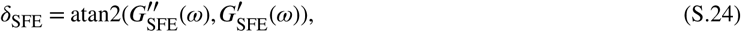

whereas the real part represents the elastic storage modulus and the imaginary part the viscous loss modulus. It is noted that *G*_0_ can be regarded as an apparent shear modulus for the time scale of *t*_0_ = 1 s.

Similar to known mechanical models, such as the Maxwell or the Kelvin-Voigt model, in which springs and dash-pots are arranged in series or parallel, the same arrangements can be applied using single fractional elements instead of simple elements. These arrangements guarantee that the equations provide mechanical and thermodynamical stability and thus are physically meaningful. ^47^ Two fractional elements can be arranged in series (Generalized Maxwell model, GM) or in parallel (Generalized Kelvin-Voigt model, GKV). The calculations for the GM is shown later in the Supplemental Information (Eq. S.35). For the GKV, the elements are arranged in parallel, meaning the stresses add and the RCE and the expression of the complex shear modulus for the GKV is

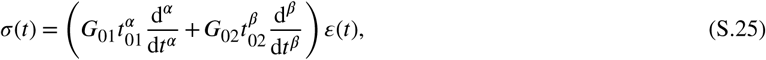

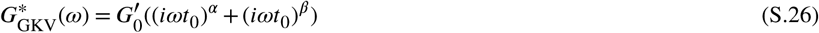

with 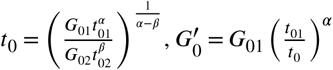 and *G*_0*i*_, *t*_0*i*_, *α* and *β* the parameters of the two fractional elements. To simplify the model and reduce the set of parameters, *G*_0_ is regarded to be independent of *α* and *β* and *t*_0_ = 1 s, and since the moduli simply add, the expression of the real and imaginary part is

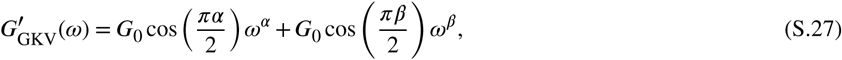

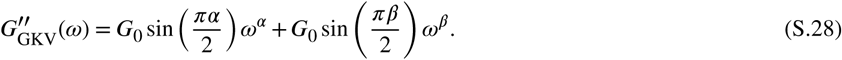

The simplification yields expressions (Eqs. S.27, S.28) with three independent parameters (*G*_0_, *α, β*). While for the SFE (Eq. S.19) there is only one power exponent *α*, the GKV shows two power exponents (*α, β*). Therefore, the GKV is able to model crossover events in which at a certain and only frequency the real part of the complex modulus equals the imaginary part. This crossover frequency for a set *t*_0_ = 1 s for the GKV is

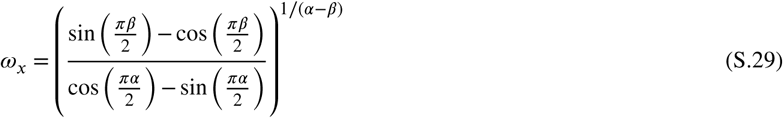

for *α* > *β* without loss of generality since *α* and *β* are interchangeable and *α* E (0.5, 1) and *β* E (0, 0.5). For the special case where *α* + *β* = 1 the crossover frequency is 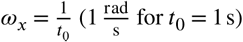. Later, in the SI (Fig. S.7) certain selected cases for different *α, β* are shown for the complex shear modulus obtained with the SFE, GM and GKV. Briefly, for frequencies *ω* ≪ *ω*_*x*_ the complex modulus behaves similar to *ω*^*β*^ for *β* > 0 and for frequencies *ω* ≫ *ω*_*x*_, *G* ∼ *ω*^*α*^. The special case of *β* = 0 shows a plateau in the real part of the complex modulus for frequencies *ω* ≪ *ω*_*x*_ while the imaginary part is ∼ *ω*^*α*^ V*ω*.

The interpretation of the GKV parameters (*G*_0_, *α, β*) are the following. *G*_0_ can be considered as an apparent shear modulus at the time scale *t*_0_ = 1 s similar to the interpretation of *G*_0_ for the SFE. However, note that *G*_GKV,0_ = 2 - *G*_SFE,0_ and is to be considered in a comparison. The power parameters *α* and *β* define the crossover frequency *ω*_*x*_ for a set *t* _0_ (see Eq. S.29). Therefore, the single information of one power parameter merely shows the complex shear modulus of the material over a certain frequency range, while the tuple (*α, β*) yields information of the complex shear modulus over the whole frequency range for a material with one crossover event.

The model using single fractional elements can be further extended by arranging multiple SFEs, such as the known mechanical models Zener model or Poynting-Thomson model with simple elements. However, this increases the amount of parameters and thus, it will not be used in this framework.

The third assumption for the microrheological analysis is dependent on the experiment. For experiments where the force is transmitted to the sample by an external bead neither the stress *σ* nor the strain *ϵ* are often directly accessible, but the force *F* acting on the bead and the bead displacement *z* instead. In the paper by Kollmannsberger *et al*. ^48^, for a magnetic tweezers experiment, the strain is estimated as the bead displacement divided by the bead radius ***R*** and the stress is estimated as the applied force divided by the bead cross section, i.e. the complex modulus 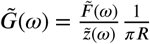. While in the paper by Balland *et al*. ^37^, for an optical tweezers experiment, the resulting complex modulus is calculated depending on the immersion of a bead inside the material characterized by an immersion half-angle *θ*, i.e.

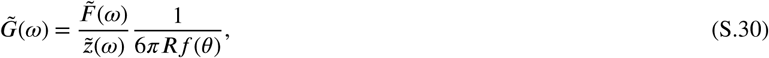

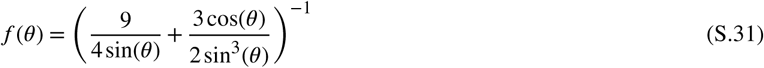

with [*f* (*θ*) *<* 1, V *θ* E (10°, 90°)], see Supp. Fig. S.9, which will be used for the analysis. For a bead immersed in an infinite medium 37,50, the complex modulus would be 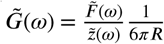.

### Generalized Maxwell Model

As mentioned, the underlying assumption is that the creep compliance behaves as a power law (Eq. S.15). In the following the results and calculations mostly based on ^47^ are summarized and presented. For the calculations of the single fractional element (SFE) and the Generalized Kelvin-Voigt (GKV) model, see the Materials and Methods section or the section on the theory of microrheology in the SI.

For the Generalized Maxwell (GM) model the fractional elements are arranged in series, meaning the individual strains add, i.e.

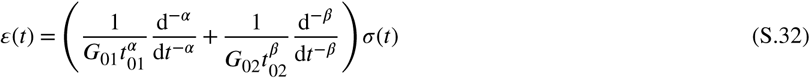

or with the fundamental relations 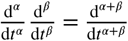

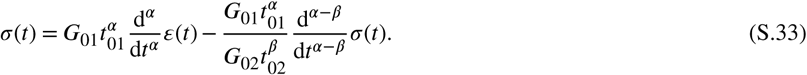

The Fourier transform yields

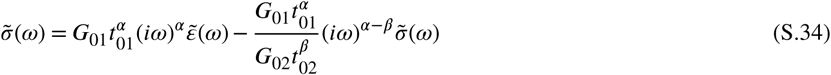

with the imaginary unit *i*^2^ = −1. With 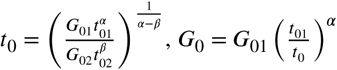 model is the complex shear modulus 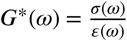 for the GM model is

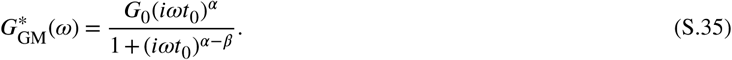

### Microrheological models using fractional calculus

As a compilation, the following equations show the complex shear modulus *G*\*(ω), including their real (*G*′(ω)) and imaginary (*G*″(ω)) part, in the Fourier domain and the creep compliance ***J*** (*t*) in time domain with 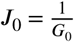 obtained from 47 for the SFE, GM and GKV. Additionally, for the GM and GKV the crossover frequency *ω*_*x*_ for *α* E (0.5, 1) and *β* E (0, 0.5) is shown. Single fractional element (SFE):

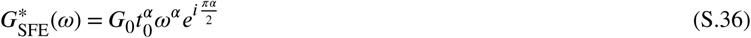

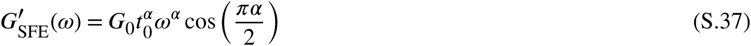

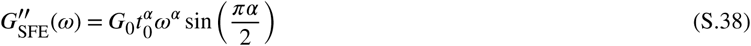

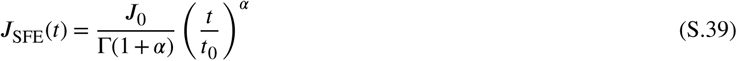

Generalized Maxwell model (GM):

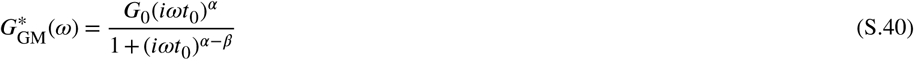

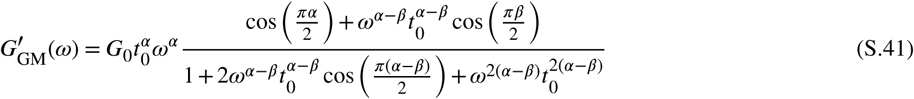

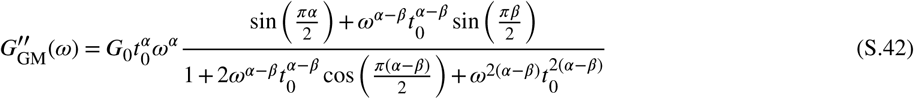

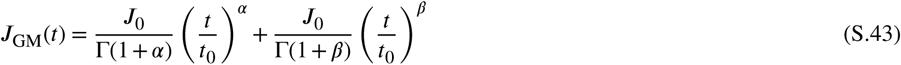

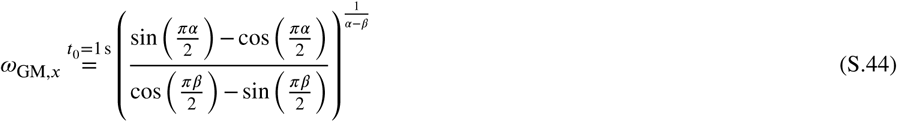

Generalized Kelvin-Voigt model (GKV):

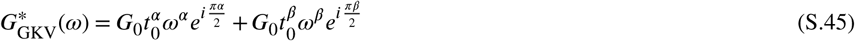

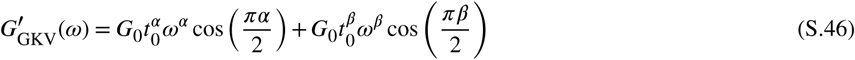

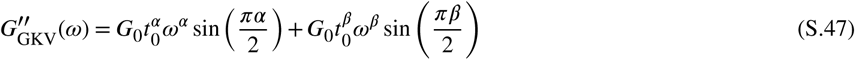

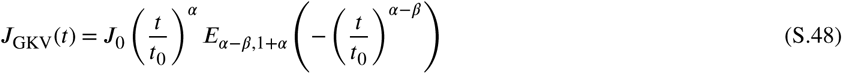

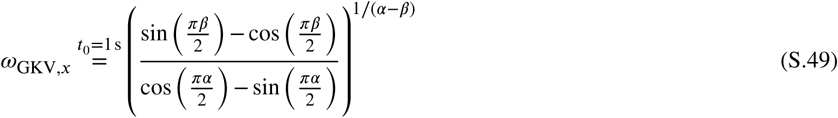

with the Mittag-Leffler function 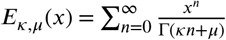.

In Supp. Fig. S.7 selected cases of the tuple (*α, β*) are shown for *G*_0_ = 1 Pa and *t*_0_ = 1 s for the GM and GKV and for the SFE the power exponent *α* is used. Without loss of generality since *α* and *β* are interchangeable, in the following the case *α* > *β* is regarded. Figure S.7A shows a case where *α* +*β* = 1 and therefore 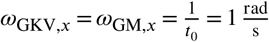. For frequencies *ω* ≪ *ω*_*x*_ the complex modulus obtained with the GM behaves as *ω*^*α*^, while with the GKV it behaves as *ω*^*β*^. Oppositely, for frequencies [*ω* ≫ *ω*_*x*_ : *G*\* ∼ *ω*^*β*^] for the GM and [*ω* ≫ *ω*_*x*_ : *G*\* ∼ *ω*^*α*^] for the GKV. Note that for *ω* < *ω*_*x*_ the imaginary part is higher than the real part for the GM and vice versa for the GKV and accordingly, for *ω* > *ω*_*x*_ the real part greater than the imaginary part for the GM and vice versa for the GKV. The complex shear modulus cannot be described with one power exponent close to the crossover frequency. Figure S.7B shows a case with *β* = 0 and *α* + *β* ≠ 1. Here, the crossover frequency is not the same for the GM and the GKV. Also, for frequencies 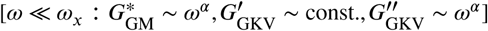 while 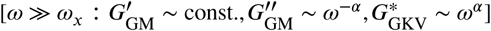. In Figure S.7C *α* = 0.2 and *β* = 0 there is no crossover frequency. Other than that, the moduli behave as in the case in Figure S.7B. For power exponents equal to zero (Fig. S.7D) the imaginary part vanishes for the SFE, GM and GKV, while the real part is constant. Note that the constant *G*_0_ is different for SFE, GM and GKV. The value of *G*_GM,0_ = 0.5 - *G*_SFE,0_ and *G*_GKV,0_ = 2 - *G*_SFE,0_, therefore to compare the absolute value of the constant *G*_0_ between the different models the scaling factor has to be considered. Figure S.7E shows the case where *α* = *β* = 0.5. In this case, the real part equals the imaginary part and they behave as *ω*^*α*^ = *ω*^*β*^. The difference between each model is due to the difference in the definition of the constant *G*_0_. In Figure S.7F the case *β* > *α* is shown with the values of *α, β* exchanged with each other compared to the case in Figure S.7A. Indeed, there is no difference in the GM and GKV model in Figure S.7A and Figure S.7F and the power exponents are interchangeable. The difference seen in the SFE model is due to the choice of *α* being the sole power exponent in the model.

Supplementary Figure S.8 shows the fit to a real data set with the different models. Due to the existence of a crossover frequency, the SFE model was used separately for the real and imaginary part of the complex shear modulus.

**Figure S.7.**
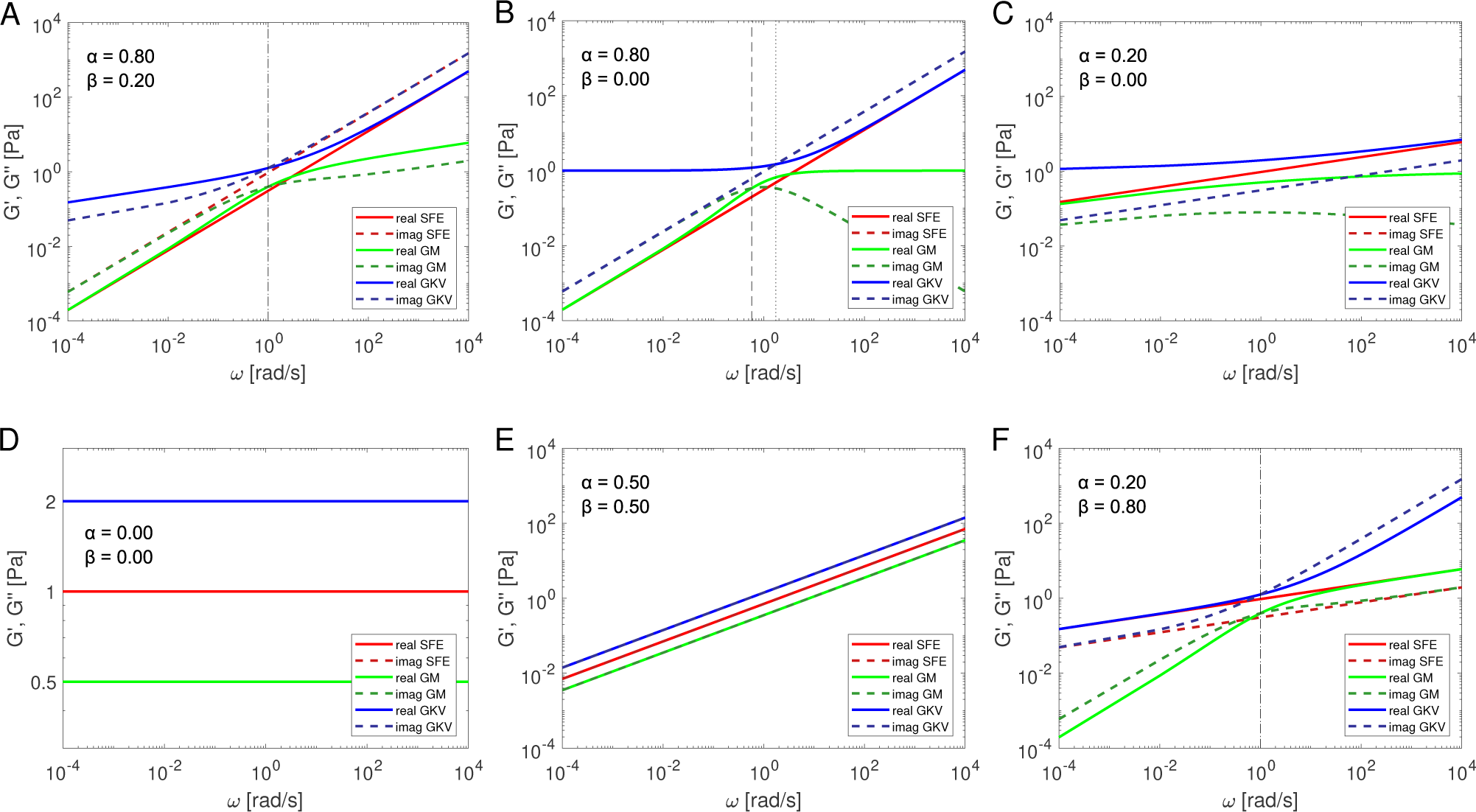
Calculated moduli for the single fractional element (SFE) model (red), Generalized Maxwell (GM) model (green) and Generalized Kelvin-Voigt (GKV) model (blue) for selected cases of the power exponents. Solid lines represent the real part of the complex shear modulus and dashed lines the imaginary part. The vertical black dashed line represents the crossover frequency of the GM model and the vertical black dotted line the crossover frequency of the GKV model. (A) Case for *α* + *β* = 1 and *α* = 0.8, *β* = 0.2. (B) Case for *α* + *β* ≠1 and *α* = 0.8, *β* = 0. (C) Case for *α <* 0.5 and *α* = 0.2, *β* = 0. (D) Case for *α* = *β* = 0. (E) Case for *α* = *β* = 0.5. (F) Case for *α < β* and *α* = 0.2, *β* = 0.8.

**Figure S.8.**
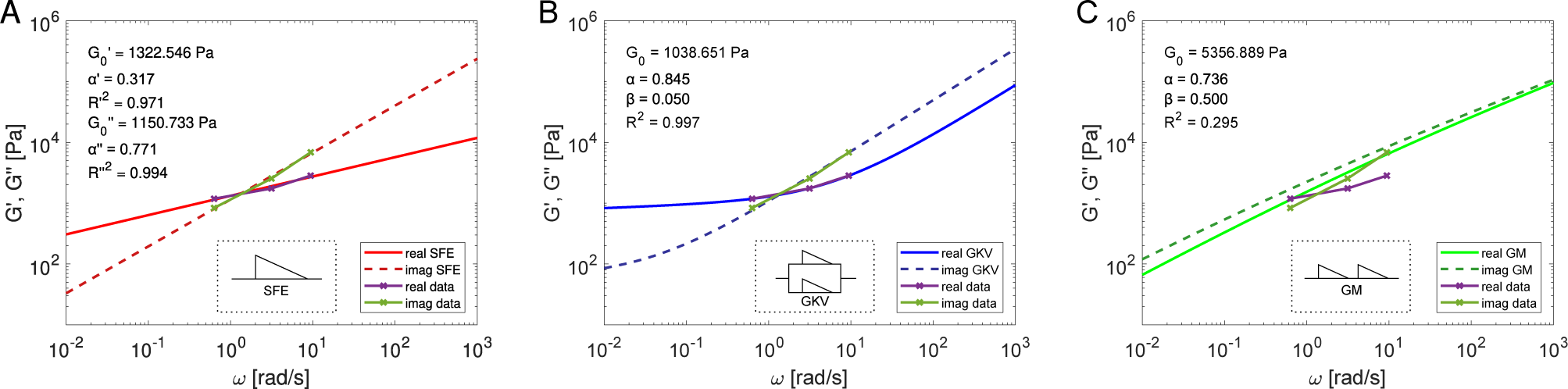
Fit of the moduli with the SFE (A, red), GKV (B, blue) and GM (C, green). Solid lines represent the real part of the complex shear modulus and dashed lines the imaginary part. For the SFE the real and imaginary part are evaluated separately. For the GKV and GM the real and imaginary part are evaluated simultaneously. Inset shows the schematic representation of the respective models after the design by Schiessel *et al*. ^47^

### Bead immersion half-angle correction

The complex shear modulus is defined as the ratio of the stress to strain. However, during the experiment only the force and the displacement of the bead with radius ***R*** is measured. Here, to estimate the stress and strain the complex modulus is calculated with a correction *f* (*θ*) based on the bead immersion half-angle *θ* into the cell

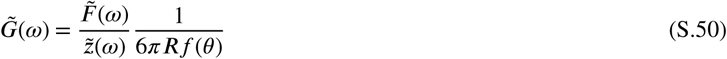

with the correction ^37^

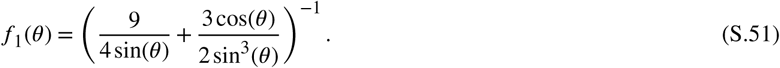

Another immersion half-angle based correction which also takes the cell height *h* at the location of the bead into consideration is given by ^49^

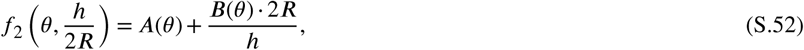

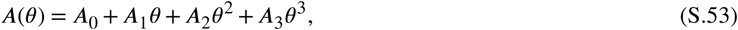

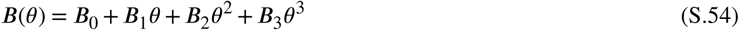

with the constant coefficients shown in Table S.8. The correction factor is also shown in Supp. Fig. S.9 for both corrections. Due to the fact that the cell height at the location of the bead was unknown during the experiment, the correction *f*_1_(*θ*) was used for the analysis. However, the height of HUVECs usually does not exceed 3 *μ*m at the cell body and is 2. 1 *μ*m at cell periphery. ^57^ The difference between the other correction 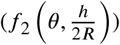 can be considered as a possible error range due to the estimation of the stress and strain.

**Table S8.** Values of constants to calculate the polynomials of the correction of the complex modulus ^49^.

| | $A_0, B_0$ | $A_1, B_1 \left[ \frac{1}{\text{rad}} \right]$ | $A_2, B_2 \left[ \frac{1}{\text{rad}^2} \right]$ | $A_3, B_3 \left[ \frac{1}{\text{rad}^3} \right]$ |
| --- | --- | --- | --- | --- |
| A | $2.321 \cdot 10^{-2}$ | $-2.054 \cdot 10^{-1}$ | $5.250 \cdot 10^{-1}$ | $-1.338 \cdot 10^{-1}$ |
| B | $4.788 \cdot 10^{-3}$ | $-4.314 \cdot 10^{-2}$ | $1.020 \cdot 10^{-1}$ | $-2.698 \cdot 10^{-2}$ |

### Dynamics of the shear modulus with the GKV model

Figure 7E shows the obtained parameter *G*_0_ using the GKV model. To test for a significant difference the obtained values are grouped into the time intervals in which the conditions had changed, e.g. a higher flow with cyto B insertion or the washing out process. The significance tests are shown in Supp. Fig. S.10A. The fit qualities with *R*^2^ > 0.7 are shown in Fig. 7E. Supplementary Figure S.10B shows the fit quality of each obtained data value. Indeed, the fit quality is mainly above *R*^2^ = 0.9, although a fit quality of *R*^2^ = 0.7 would still be valid for a simultaneous fit of the real and imaginary part of the complex shear modulus.

**Figure S.9.**
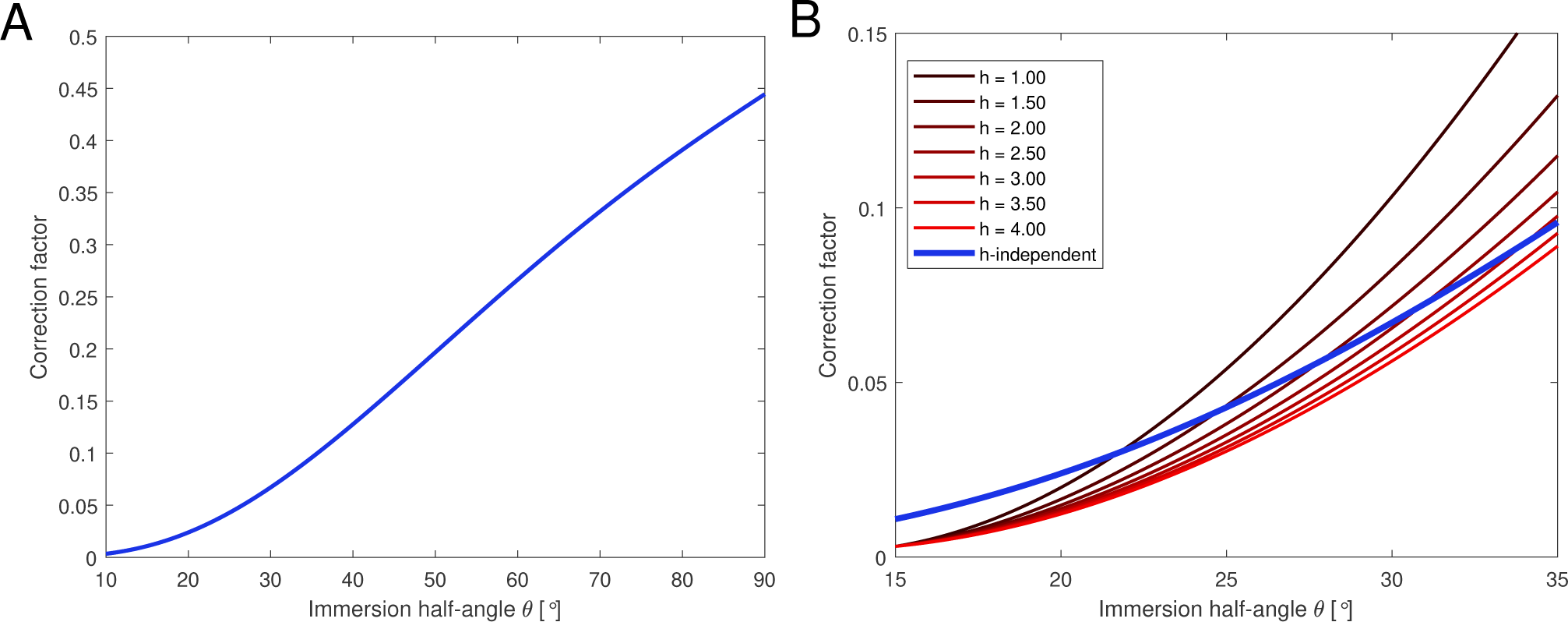
Different correction factors to calculate the complex modulus with Eq. S.5. (A) Correction factor *f*_1_(*θ*) in the range of [10°, 90°] obtained from ^37^ which is also used in the analysis. (B) Correction factor 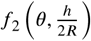 obtained from ^49^ which also takes the cell height *h* into consideration as indicated in red colors and the correction factor *f*_1_(*θ*) (blue).

**Figure S1.0.**
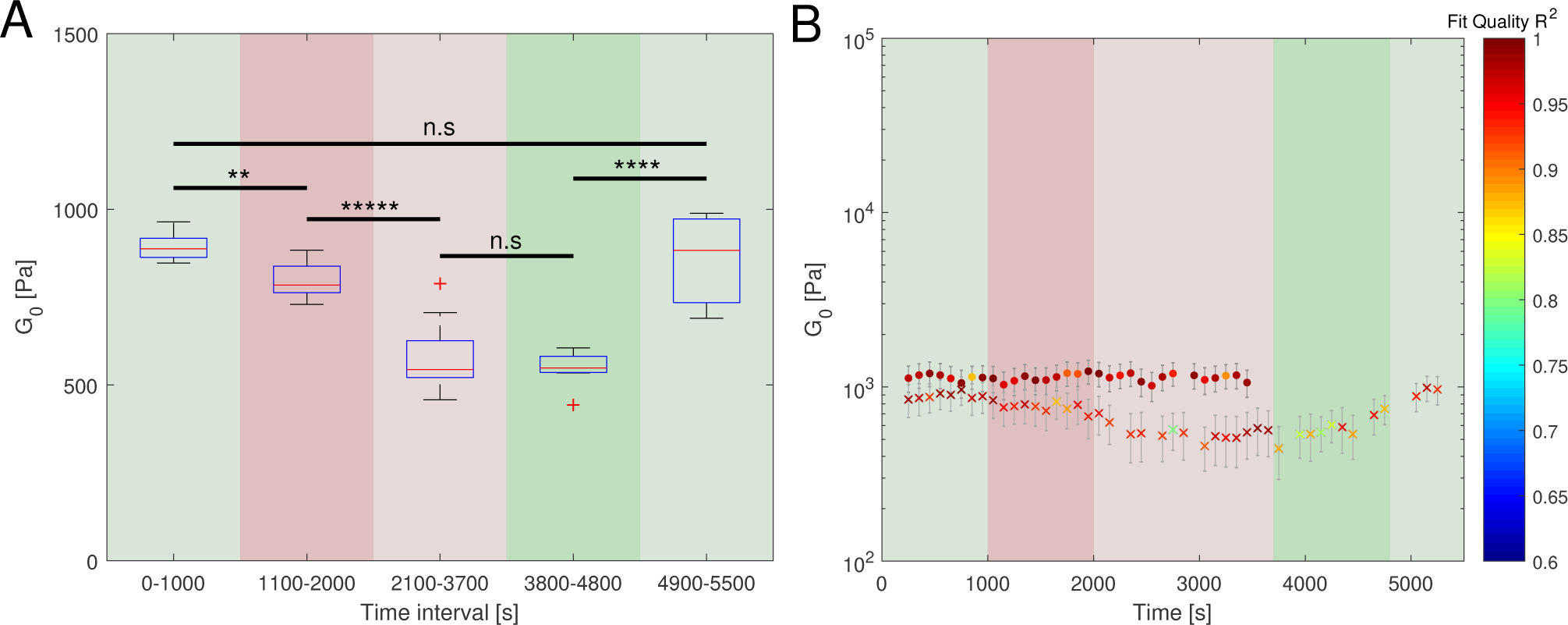
(A) Significance tests between different time intervals and conditions for the values of *G*_0_ obtained from the GKV model for the cyto B experiment. The significance star * represents *p* ≤ 0.05, ^**^ : *p* ≤ 0.01, ^***^ : *p* ≤ 0.001, ^****^ : *p* ≤ 0.0001, ^*****^ : *p* ≤ 0.00001 and n.s : not significant, using a two-sample *t*-test. (B) Representation of Figure 7E with the fit quality *R*^2^ for each obtained data value.

### Data post-processing and analysis

The relevant, measured data are the positional data in *x, y, z*, the amplitude corresponding to the exerted acoustic force *F* and the sampling time *t*. A complete data set consists of the measured data in the full duration of acquisition, e.g. a duration of 3700 s. The following shows the automatized steps of the analysis using the self-written MATLAB software *Kitsune (version 0*.*98)*.

- Find the start and end time of the mOsc corresponding to the data with force type *F*_*t*_ = 5 in the modified LabVIEW software. (Full force span)
- Segment the full force span in sections of 500 s evaluation time intervals in 100 s time steps.
- Analyze each of the sections separately with the following steps.
- Get the mean *x, y*-position during the analyzed section and calculate the force value according to the conversion factor from the calibration map.
- Subtract the force values of the section with its mean to avoid the errors from the zero-padding in a later step. This is valid because the oscillation is of interest and not the initial force step.
- Clean the *z*-data from tracking errors, e.g. due to particles interfering with the tracking’s region of interest (RoI) during flow.
  - Segment the section into 1 s sections. (Error-Section)
  - Calculate the median of the Error-Section. (Local-Median)
  - Find data points within the Error-Section and replace them with the Local-Median if their difference with the Local-Median is greater than 500 nm.
  - Calculate the median of the whole section. (Global-Median)
  - Find data points within the Error-Section and replace them with the Global-Median if their difference with the Global-Median is greater than 10000 nm.
- Correct the *z*-drift using the continuous poly2 subtraction. (Note that it is not necessarily smooth, but only continuous.)
  - Segment the section into 25 s sections and the final section with at least 3 data points else it is filled with the last value. (Drift-Sections)
  - Create the Vandermonde Matrix for a polynomial of 2nd order (*n* = 2).
  - Constraint the fit: The starting point has to be the ending point of the previous Drift-Section; for the first Drift-Section the starting point of the data is also its starting point.
  - Fit the data using lsqlin and polyval which are in-built functions of MATLAB.
  - Append all Drift-Sections to a data set. (Drift-Correction)
  - Subtract the section with the Drift-Correction.
- Filter out sections (after the correction from above) that have high errors, e.g. a particle interfered with the RoI for too long or the tracked particle is no longer being tracked.
  - Calculate the standard deviation (std) of the *z*-position in the section.
  - Set the results to NaN and stop the analysis if at least one of the following criteria are fulfilled:
    1. std(*z*) > 1000 nm
    2. std(*z*) == 0
    3. abs(median(*z*)) *<*= 10^−8^ nm
    4. abs(median(abs(*z*)) −mean(abs(*z*))) > 200 nm
    5. abs(median(*z*)) > 1000 nm
    6. abs(mean(*z*)) > 1000 nm
    7. abs(abs(mean(abs(*z*(1st half)))) −abs(mean(abs(*z*(2nd half))))) > 50 nm
- Calculate the Discrete-Time Fourier Transformation (DTFT) of *z* and *F*.
  - Calculate the mean time steps between each data point inside the section due to possible sampling time errors.
  - From the mean time step, calculate the mean sampling frequency.
  - If the length of the data points inside the section is odd, remove the last data point to make it an even length.
  - Calculate the DTFT by zero-padding the data until the length is 100-times the data length.
  - Multiply the DTFT by the mean time step and by 2 for the normalization and due to the fact of considering the positive frequency only.
- Find the peaks of the absolute of the DTFT.
  - Get the maximum of the DTFT within a 0.01 Hz interval around the estimated driving frequencies.
- Calculate the complex modulus *G*\* using Eq. S.50 with the peaks of the DTFT of z and the peaks of the DTFT of the force and the determined angle, here *0* = 28.73°.
- The mean absolute, real and imaginary part of *G*\* of all *n* beads is calculated with the standard error 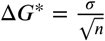 and the standard deviation *σ*.
- The errors of the ratio of real and imaginary part as well as the phase is calculated with error propagation.
- The fit parameters (*G*_0_, *α*) of the SFE (power) model is obtained by fitting to the logarithm of the Eq. S.19 for real and imaginary part separately by minimizing the value 1 − *R*^2^ for each fit, respectively. The logarithmic value of *G*_0_ is transformed appropriately. The error is obtained with *n*_boot_ = 100 bootstrap samples. The values are set to NaN if *R*^2^ *<* 0.9.
- The fit parameters (*G*_0_, *α, β*) of the GKV model is obtained by fitting the real part of the complex shear modulus to the real part of Eq. S.26 and the imaginary part to the imaginary part of Eq. S.26. The sum of 1 − *R*^2^ of each fit is then minimized. The error is obtained with *n*_boot_ = 100 bootstrap samples. The values are set to NaN if *R*^2^ *<* 0.7.

### SFC Protocol

This protocol is used to perform the Stokes Force Calibration (SFC) on Acoustic Force Spectroscopy (AFS) chips. It also serves as a guide. It is part of the Supplemental Information of the publication *Microchip based microrheology via Acoustic Force Spectroscopy shows that endothelial cell mechanics follows a fractional viscoelastic model*. The calibration method is described in the main text of the publication and the recommended software *Kitsune* to analyze the SFC data is available on github https://github.com/A141-User/Acoustic-Kitsune. This protocol is for the G2 AFS chips; other generations may differ.

### Conditions of the main experiment

The conditions of the main experiment has to be set beforehand. Firstly, to ensure that the AFS can perform under the conditions and, secondly, to perform the SFC under the same conditions due to the dependency of the force on the respective conditions.

Conditions:

- Temperature (*T <* 40°)
- Medium (liquid during experiment)
- Bead size (preferably 0.5 *μ*m *< d <* 20 *μ*m)
- Bead type (e.g. polystyrene, silica, etc.)
- Desired force range (pN-nN)
- (optional) to think about:
  - force type, e.g. constant force, force ramp, etc.
  - length of force application

### Preparation

Prior to a successful calibration there are still a few aspects to be taken care of.

- This calibration method is based on video-capturing, meaning that a camera with a sampling rate of *f*_*s*_ > 30 Hz is recommended.
- The surface of the AFS fluid chamber is made of glass. The beads in combination of the medium may bind to the surface, e.g. due to charge. Therefore, the glass or the beads can be coated accordingly to prevent specific binding on the surface.
- The bottom of the chip should be cleaned to prevent dirt that can disturb the tracking and may also alter the force values.
- The inside of the fluid chamber should be cleaned, e.g. with ethanol or standard bleach (e.g. A1727 Sigma-Aldrich) and thoroughly washed out afterwards.
- The position of the AFS chip is fixed and can only be moved in a controlled manner to ensure to measure at the same field of view (FoV).

### Procedure

After meeting the conditions of the main experiments and a successful preparation the SFC can be performed. The temperature, medium and beads mentioned here are the ones of the main experiment for which the following calibration is performed.

1. set the temperature (wait at least 1 min)
2. suspend the beads in the medium (bead suspension)
3. insert the bead suspension into the chip
4. let the beads fall to the ground by gravity

Prior to actually measuring the beads, it might be more feasible to just observe the beads in the bright-field image when applying a low amplitude first to quickly scan through the chip for usable FoVs. Find the desired FoV and make sure to be able to find the same FoV even after unmounting the chip from the microscope.

It is recommended to use the resonance frequency because the ratio of the axial force to the lateral force is typically higher at the resonance frequency. The following steps are to obtain the resonance frequency.

1. track all single beads
2. create a look-up-table (LUT) in the range greater than the first *z*-node, typically 20 *μ*m for a specific type of chips
3. measure all beads (here, if possible with a moving region of interest (RoI))
4. as a first estimation, use the recommended frequency saved inside the chip
5. apply a constant pressure amplitude for 1 s
6. change the frequency by fl*f* = 0.01 MHz (in both directions)
7. after the bead is at the ground again, repeat the same constant pressure amplitude for 1 s
8. repeat steps 6. and 7. until the resonance frequency is found which can be seen by the measured *z*-position (fastest rise to the *z*-node) or a quick analysis using *Kitsune* (highest force)
9. to prepare the measurement to obtain the conversion factor which requires several constant pressure amplitude values, check for the highest possible amplitude at which the *z*-position of the bead can be sufficiently recorded by the camera

Now, the resonance frequency is found and the SFC can be performed to obtain the spatial calibration map of the FoV showing the conversion factors using the following steps.

1. insert new beads
2. wait until they are on the ground
3. track, create LUT and measure the positional data of the beads (do not use a moving RoI)
4. apply a low amplitude for 1 s
5. wait until they are on the ground
6. apply a higher amplitude for 1 s
7. repeat steps 5. and 6. until a desired amount of amplitudes are measured (preferably 4 different amplitudes smaller than the highest possible to capture)
8. end measurement
9. repeat steps 1-8 until a desired amount of beads are measured, i.e. the measured beads are well-distributed throughout the whole FoV (which can be quickly checked using *Kitsune*’s “Bead distribution”)

The experimental procedure is completed and the *xyz*-positional data of every bead has been measured. For every bead, the *z*-position has to be evaluated during the force application. The lateral location of each bead is known and the conversion factor is assigned to each bead to obtain the distribution of the conversion factor. In order to obtain the calibration map of the conversion factors, the distribution has to be interpolated. The analysis can be performed using the steps shown in the main text of the publication or using *Kitsune*.

### Further steps

After obtaining the calibration map for one FoV at one specific condition, the calibration is basically successful. A few other steps can be performed, as listed in the following.

- find other FoVs and obtain the maps (calibration), make sure to assign the maps to the FoV accordingly
- calibrate the FoV at different conditions, such as different temperatures
- measure the temperature change inside the flow chamber during the SFC force with an external temperature sensor if the temperature sensor inside the chip holder cannot measure the temperature inside the flow chamber
- measure the temperature change inside the flow chamber during force application of the main experiment
- obtain the temperature dependency of the force at the FoV if the temperature change is significantly high
- perform the main experiment
- calibrate the same FoV at the same condition again after a while to account for degradation
- calibrate other AFS chips

### Additional notes

- The analysis software *Kitsune* is available on github https://github.com/A141-User/Acoustic-Kitsune and is based on Matlab R2017 (The MathWorks). It is written for data generated by a modified version of the Generation 2 AFS LabVIEW tracking software. However, it can also analyze the standard version.
- The density of the bead has to be greater than the density of the medium under the conditions.
- If a force type other than the constant force or force ramp is needed, the LabVIEW software may have to be modified.
- If a high force amplitude is applied, the change of the temperature inside the fluid chamber during force application should be measured.

